# Targeting Cardiac Fibrosis with Chimeric Antigen Receptor Neutrophils Derived from Human Pluripotent Stem Cells

**DOI:** 10.64898/2026.10.11.758383

**Authors:** Gyuhyung Jin, Shaohai Fang, Yun Chang, Kunming Shao, Huiyang Li, Yan Tan, Xiaojun Lance Lian, Qing Deng, Yun Nancy Huang, Xiaoping Bao

## Abstract

Heart failure remains a leading cause of global morbidity and mortality, with cardiac fibrosis contributing to disease progression. Following myocardial injury, activated cardiac fibroblasts orchestrate adverse extracellular matrix (ECM) remodeling, which leads to tissue stiffening and compromised contractility. Despite the clinical importance of cardiac fibrosis, current pharmacological interventions modulate disease progression without directly eliminating activated fibrogenic cells. To target these cells directly, we engineered human pluripotent stem cell (hPSC)-derived neutrophils expressing a chimeric antigen receptor (CAR) directed against fibroblast activation protein (FAP). In vitro, FAP CAR neutrophils exhibited increased reactive oxygen species-associated fluorescence during fibroblast co-culture and dose-dependent cytotoxicity in 2D cultures, while reducing fibroblast-associated fluorescence in 3D cardiac organoids without a significant reduction in cardiomyocyte-associated cardiac troponin T staining. FAP CAR neutrophils also showed increased iNOS and decreased Arginase-1 expression compared with unmodified neutrophils, consistent with a shift toward an N1-associated polarization profile. Analysis of public cardiac single-cell and single-nucleus RNA-sequencing datasets identified temporal patterns of immune-associated expression and fibroblast activation that informed treatment timing. In a murine myocardial infarction (MI) model, mice treated systemically with FAP CAR neutrophils showed higher day 7 left ventricular ejection fraction than PBS-treated mice in an exploratory comparison. Together, these findings support further evaluation of hPSC-derived FAP CAR neutrophils as an immunotherapeutic approach to cardiac fibrosis.

## 1. Introduction

Cardiac fibrosis is a common pathological feature observed in many forms of heart disease, accounting for substantial morbidity and mortality globally [1]. The central histopathologic feature of fibrotic cardiac tissue is the aberrant accumulation and remodeling of extracellular matrix (ECM) proteins, predominantly secreted by activated cardiac fibroblasts, or myofibroblasts [2]. Although deposition of structural proteins such as collagen initially helps prevent ventricular rupture after injury, persistent deposition impairs myocardial compliance, disrupts electrical conduction, and contributes to adverse outcomes such as cardiomyocyte hypertrophy, arrhythmias, and heart failure. Current clinical management relies heavily on pharmacological agents that block upstream signaling pathways involved in fibroblast activation, including the renin-angiotensin-aldosterone system [3]. While these therapies can slow disease progression, they do not directly eliminate existing fibrogenic cells, leaving established ECM remodeling incompletely addressed [4]. These limitations motivate approaches that directly target pathogenic fibrogenic cells to limit fibrosis and promote more favorable remodeling.

Recent studies in adoptive cellular immunotherapy have shown that pathogenic stromal cells can be targeted directly. Chimeric antigen receptor (CAR) T cells engineered to target fibroblast activation protein (FAP), a stromal marker upregulated on activated cardiac fibroblasts, have demonstrated significant antifibrotic efficacy and functional restoration in murine models of cardiac injury [5,6]. Despite these encouraging findings, translating T cell–directed strategies to non-malignant indications presents several challenges. Ex vivo CAR-T approaches involve manufacturing complexity and concerns related to long-term lymphoid persistence [7,8]. In vivo mRNA-LNP platforms present different challenges, including systemic biodistribution, variable transduction efficiency, and immunogenicity associated with repeat dosing [9].

Innate immune cells have also been explored as alternative effectors for targeting cardiac fibrosis. Strategies such as FAP-targeted CAR-macrophages [10,11] and CAR-monocytes [12] offer tissue-infiltrating and phagocytic effector functions for targeting myofibroblasts, while engineered tolerogenic dendritic cells (DCs) [13] have shown potential in modulating the immune response via regulatory T cell expansion. Yet, the phenotypic plasticity of macrophages and monocytes renders them susceptible to the local cytokine milieu; in the injured heart, signals associated with alternative macrophage activation may also support fibroblast activity and matrix remodeling [14–16]. Tolerogenic DC approaches also depend on indirect immune modulation and maintenance of a regulatory phenotype [17], highlighting the value of evaluating additional effectors capable of rapid, direct targeting with a shorter duration of activity.

Given these limitations, we investigated neutrophils as an alternative effector-cell population. Neutrophils possess a capacity for rapid extravasation, allowing them to reach injured tissues and participate in local remodeling [18]. Their short lifespan may limit prolonged effector exposure, making them candidates for transient cell therapy [19]. Because primary neutrophils are difficult to expand and genetically modify ex vivo [20], we utilized a renewable, scalable human pluripotent stem cell (hPSC) platform. Generating CAR neutrophils from a renewable hPSC source provides a route toward standardized, bankable products while enabling CRISPR-mediated genetic engineering at the stem cell stage.

Using our previously reported differentiation platform for producing tumor-targeting CAR neutrophils [21,22], we hypothesized that hPSC-derived neutrophils (hNeu) engineered with a FAP-directed CAR would selectively eliminate activated cardiac myofibroblasts and attenuate myocardial fibrosis without compromising healthy tissue. Given the liabilities associated with dysregulated granulocyte activation, including indiscriminate degranulation, reactive oxygen species (ROS) release, and bystander tissue damage [23,24], we examined target-associated activation and fibroblast-directed activity alongside potential effects on neighboring cardiomyocytes. In this study, we generated FAP CAR neutrophils from hPSCs (hNeu-FAP) and established an intermediate progenitor cryopreservation protocol to support cell banking before neutrophil differentiation. Bulk RNA sequencing identified differences in inflammatory and matrix-remodeling transcripts in CAR-engineered neutrophils, and flow cytometry showed increased iNOS and decreased Arginase-1 expression in hNeu-FAP. We then evaluated fibroblast-targeting activity in 2D co-cultures and 3D human cardiac organoid models. In these models, hNeu-FAP showed activity against activated epicardial fibroblasts, and treated organoids showed no statistically significant reduction in cardiomyocyte-associated cTnT fluorescence relative to untreated controls. Analysis of temporal immune- and fibroblast-associated expression across cardiac single-cell datasets informed the selection of an early post-injury intervention time point. Finally, we assessed the presence of transferred hNeu in ISO-injured myocardium and evaluated left ventricular ejection fraction and scar area after FAP CAR neutrophil administration in a separate murine myocardial infarction model. Together, these findings support a progenitor-banking strategy and the further development of FAP-directed neutrophil immunotherapy for cardiac fibrosis.

## 2. Materials and Methods

### 2.1 hPSC Culture and Genome Engineering

H9 human embryonic stem cells (WiCell) were maintained on Matrigel-coated plates in mTeSR Plus medium under standard conditions. To generate FAP-targeting CAR-hPSCs, cells were nucleofected with an AAVS1-targeting CRISPR/Cas9 ribonucleoprotein/plasmid complex alongside a donor template encoding an anti-FAP scFv CAR construct. Transfected clones were enriched under puromycin selection, isolated, and verified for targeted genomic integration via PCR genotyping.

### 2.2 Directed Lineage Differentiation and Organoid Assembly

For lineage differentiation, hPSCs were directed toward hematopoietic and neutrophil lineages via stage-specific modulation of Wnt signaling and sequential treatment with hematopoietic cytokines, including VEGF, SCF, FLT3L, GM-CSF, IL-3, and G-CSF, with progenitor cells and mature neutrophils harvested or cryopreserved at defined developmental stages. In parallel, cardiac progenitors and downstream lineages, including epicardial-derived fibroblasts and cardiomyocytes, were generated through biphasic Wnt pathway modulation, followed by TGF-β–mediated activation and lentiviral transduction with GFP or luciferase reporters where indicated. Three-dimensional cardiac fibrosis organoids (hCFOs) were subsequently self-assembled by co-culturing cardiomyocytes and activated epicardial fibroblasts in non-adherent U-bottom plates, into which engineered or control neutrophils were introduced to evaluate targeted cytotoxicity and anti-fibrotic activity.

### 2.3 Functional, Cytotoxicity, and Migration Assays

Target cell killing by CAR-neutrophils was quantified using bioluminescence-based lysis assays with luciferase-expressing target fibroblasts at indicated effector-to-target (E:T) ratios, while cell viability was assessed using the CellTiter-Glo luminescent assay. Neutrophil chemotaxis and directional migration were evaluated under agarose overlays using time-lapse microscopy, and antigen-specific binding alongside phagocytic activity were determined using fluorophore-conjugated recombinant FAP and bioparticle conjugates. Additionally, ROS generation following chemical or target-cell stimulation was quantified using fluorogenic oxidative stress indicators on a microplate reader.

### 2.4 Flow Cytometry and Immunofluorescence

Surface marker expression, intracellular lineage markers, and cell purity were determined using standard flow cytometry protocols on an Accuri C6 Plus platform. For structural evaluation, 2D cultures and intact 3D organoids were fixed with 4% PFA, permeabilized, and stained with validated primary and fluorophore-conjugated secondary antibodies. Confocal (Zeiss LSM900) and fluorescence imaging were performed to analyze cell spatial distribution, immune synapse formation, and tissue architecture. Image analysis, radial profiling, and 3D surface mapping were conducted using ImageJ/Fiji and Origin software.

### 2.5 Transcriptomic Profiling and Bioinformatic Analysis

Total RNA from control and CAR-neutrophils was sequenced, and differential gene expression was evaluated using DESeq2 in R, defining significant DEGs under Benjamini-Hochberg FDR thresholds (adjusted P < 0.05, |Log2FC| > 1). For single-cell transcriptomics, publicly available murine cardiac injury datasets (GEO and ArrayExpress accessions) were processed using Scanpy (v1.9+) in Python; following standard quality control filtering and normalization, cellular identities across the cardiac cellulome were resolved using curated canonical marker gene modules and visualized via UMAP embeddings and dot plots.

### 2.6 In Vivo Studies and Myocardial Delivery

All animal procedures were approved by the respective Institutional Animal Care and Use Committees (IACUC) at the participating institutions. Acute myocardial injury was induced in mice via isoproterenol administration to assess neutrophil homing; endogenous leukocyte infiltration and the recruitment of adoptively transferred human neutrophils were evaluated in transcardially perfused hearts by flow cytometric quantification of tissue-digested single cells and immunohistological evaluation of ventricular cryosections. To assess therapeutic efficacy in ischemic injury, myocardial infarction (MI) was surgically induced in immunodeficient mice via permanent coronary artery ligation. Engineered FAP-CAR or unmodified hPSC-derived neutrophils, or vehicle control, were adoptively transferred post-injury. Cardiac function was longitudinally monitored using echocardiography, and myocardial remodeling alongside scar area resolution was evaluated by histological staining of ventricular tissue sections.

### 2.7 Statistical Analysis

Data are presented as mean ± SD. Statistical comparisons between two groups were evaluated using two-tailed unpaired Student’s t-tests. Multi-group and longitudinal comparisons were determined by one-way or two-way ANOVA followed by Tukey’s post hoc test in GraphPad Prism. Significance thresholds are designated in the respective figure legends.

Note: Detailed step-by-step protocols, primer sequences, and specific reagent catalog numbers are available upon peer-reviewed publication or from the corresponding author upon reasonable request.

## 3. Results

### 3.1 Generation and Characterization of FAP CAR hPSCs and Differentiation to Neutrophils

To specifically target activated cardiac fibroblasts, we genetically engineered hPSCs with a CAR directed against FAP. A single-chain variable fragment (scFv, clone MO36) exhibiting cross-reactivity toward both human and murine FAP [26] was synthesized and cloned into an AAVS1 donor targeting vector driven by a constitutive CAG promoter. Flanked by left and right AAVS1 homology arms, the FAP CAR expression cassette was integrated into the AAVS1 safe-harbor locus via CRISPR/Cas9-mediated homology-directed repair (HDR) (**Supplementary Figure 1A**). Junction-PCR genotyping of isolated single-cell clones supported targeted genomic integration, identifying multiple monoallelic (heterozygous) and biallelic (homozygous) knock-in lines (**Supplementary Figure 1B**). A homozygous clone was selected, expanded, and used for downstream studies.

Both unmodified and FAP CAR hPSCs maintained typical colony morphology and expression of OCT4 and NANOG, as confirmed by immunofluorescence microscopy and flow cytometry (**Supplementary Figure 1C–E**). Antigen association with the engineered cells was assessed by overnight incubation with recombinant FITC-conjugated FAP (FAP-FITC). Subsequent live-cell fluorescence imaging and quantitative flow cytometry demonstrated substantially greater cell-associated FAP-FITC fluorescence in the engineered FAP CAR hPSCs than in unmodified controls (**Supplementary Figure 1F–H**).

To determine whether stable CAR integration influences hematopoietic lineage specification, unmodified and FAP CAR hPSCs were subjected to a defined, stage-specific differentiation protocol toward neutrophils [21] (**Figure 1A**). Monolayer hPSCs initially transitioned through a hemogenic endothelial intermediate that continuously released non-adherent human hematopoietic progenitor cells (hHPCs) positive for pan-hematopoietic markers CD43 and CD45 (**Figure 1B**). Flow cytometric profiling at day 15 of differentiation revealed that approximately 80% of the emerged suspension cells co-expressed CD43 and CD45 in both unmodified and engineered lines (**Figure 1C, D**).

**Figure 1.**
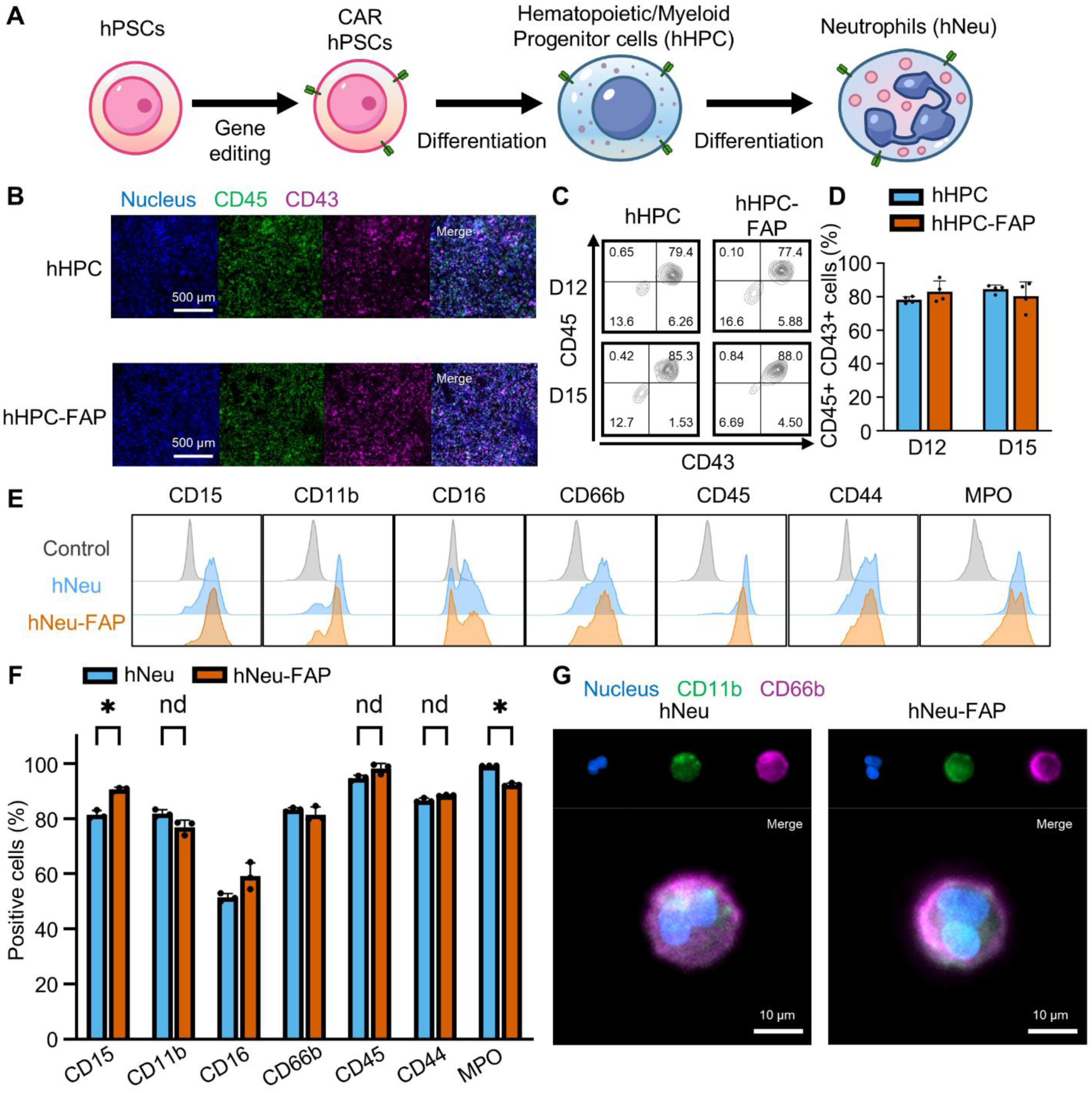
Differentiation of CAR hPSCs into CAR neutrophils. (A) Schematic overview of CAR gene editing in human pluripotent stem cells (hPSCs) and subsequent differentiation into hPSC-derived hematopoietic/myeloid progenitor cells (hHPCs) and neutrophils (hNeu). (B) Representative immunofluorescence (IF) images of wild-type hHPCs and FAP CAR hHPCs (hHPC-FAP) stained for pan-hematopoietic progenitor markers CD43 and CD45 at day 12 of differentiation. Scale bars, 500 µm. (C) Representative flow cytometry contour plots for CD43 and CD45 in floating hHPCs and hHPC-FAPs at day 12 (D12) and day 15 (D15) of neutrophil differentiation, with (D) corresponding quantification. (E) Representative flow cytometry histograms of neutrophil and myeloid maturation markers (CD15, CD11b, CD16, CD66b, CD45, CD44, and MPO) in hNeu and FAP CAR hNeu (hNeu-FAP) at day 22 of differentiation, with (F) corresponding quantification. (G) Representative IF images demonstrating CD11b and CD66b expression in hNeu and hNeu-FAP. Scale bars, 10 µm. Data are represented as mean ± SD. Statistical significance was determined by unpaired Student’s t-tests comparing hNeu vs. hNeu-FAP (n = 3 biological replicates). *P < 0.05.

Harvested day 15 hHPCs were subsequently re-plated in myeloid-instructive conditions for further neutrophil differentiation. Both hNeu and hNeu-FAP expressed signature myeloid and neutrophil surface antigens including CD11b, CD15, CD16, CD44, CD45, and CD66b along with intracellular myeloperoxidase (MPO), with broadly similar marker-positive fractions but detectable differences in CD15 and MPO (**Figure 1E, F**). Immunofluorescence confirmed CD11b and CD66b expression together with the characteristic multilobed nuclear morphology (**Figure 1G**). Collectively, these findings indicate that the FAP CAR-engineered line retains pluripotency-marker expression and the capacity for directed differentiation into neutrophils.

### 3.2 Cryopreservation and Post-Thaw Differentiation of Neutrophil Progenitors

A practical off-the-shelf workflow requires a reliable cryopreservation strategy. Because primary mature neutrophils are short-lived and difficult to cryopreserve [27,28], we first evaluated the post-thaw viability of terminally differentiated hNeu. Following cryopreservation in DMSO/serum-containing medium, thawed hNeu were tracked over time using Calcein AM staining (**Supplementary Figure 2A**). Post-thaw viability decreased substantially within the first 24 hours, with near-total cell loss observed by day 2, indicating poor recovery of mature hNeu under the cryopreservation conditions tested. To bypass this limitation, we investigated whether cryopreservation could instead be performed at an intermediate progenitor stage without compromising downstream differentiation capacity. Cells were cryopreserved at day 12 (D12) and thawed to test maturation potential under different cytokine regimens (**Supplementary Figure 2B**). Post-thaw supplementation with either GM-CSF or G-CSF from D12 to D15, followed by G-CSF terminal differentiation through D20, yielded comparable frequencies of CD16⁺ CD66b⁺ neutrophils (**Supplementary Figure 2B**).

We next compared cryopreservation timing (D12 vs. D15) and evaluated whether human serum supplementation during post-thaw maturation impacts differentiation efficiency and overall viability (**Supplementary Figure 2C–E**). Flow cytometric analysis of neutrophil maturation markers CD66b and CD11b revealed that progenitors frozen at D12 generated significantly higher proportions of CD66b⁺ CD11b⁺ double-positive cells compared to those cryopreserved at D15 (P < 0.001), while the presence of human serum did not significantly alter maturation marker expression (**Supplementary Figure 2C, D**). However, assessment of post-thaw metabolic viability by CellTiter-Glo assay demonstrated that human serum supplementation significantly increased ATP-associated luminescence across both cohorts, with D12-cryopreserved progenitors yielding higher signals than D15 progenitors, consistent with improved viable-cell recovery (P < 0.01; **Supplementary Figure 2E**). Based on these results, we selected D12 myeloid progenitor cryopreservation followed by human serum-supplemented maturation as the preferred protocol among the conditions tested.

### 3.3 Molecular and Functional Characterization of hNeu and hNeu-FAP

To determine whether CAR engineering alters basic neutrophil physiology, we performed morphological and functional characterization of the differentiated cells, together with transcriptomic analysis of CAR-engineered neutrophils. Wright-Giemsa staining demonstrated characteristic polymorphonuclear, multi-lobed nuclear architecture in both hNeu and hNeu-FAP (**Figure 2A**), a structural feature essential for cellular deformability and transmigration through confined microvascular and interstitial barriers [29].

**Figure 2.**
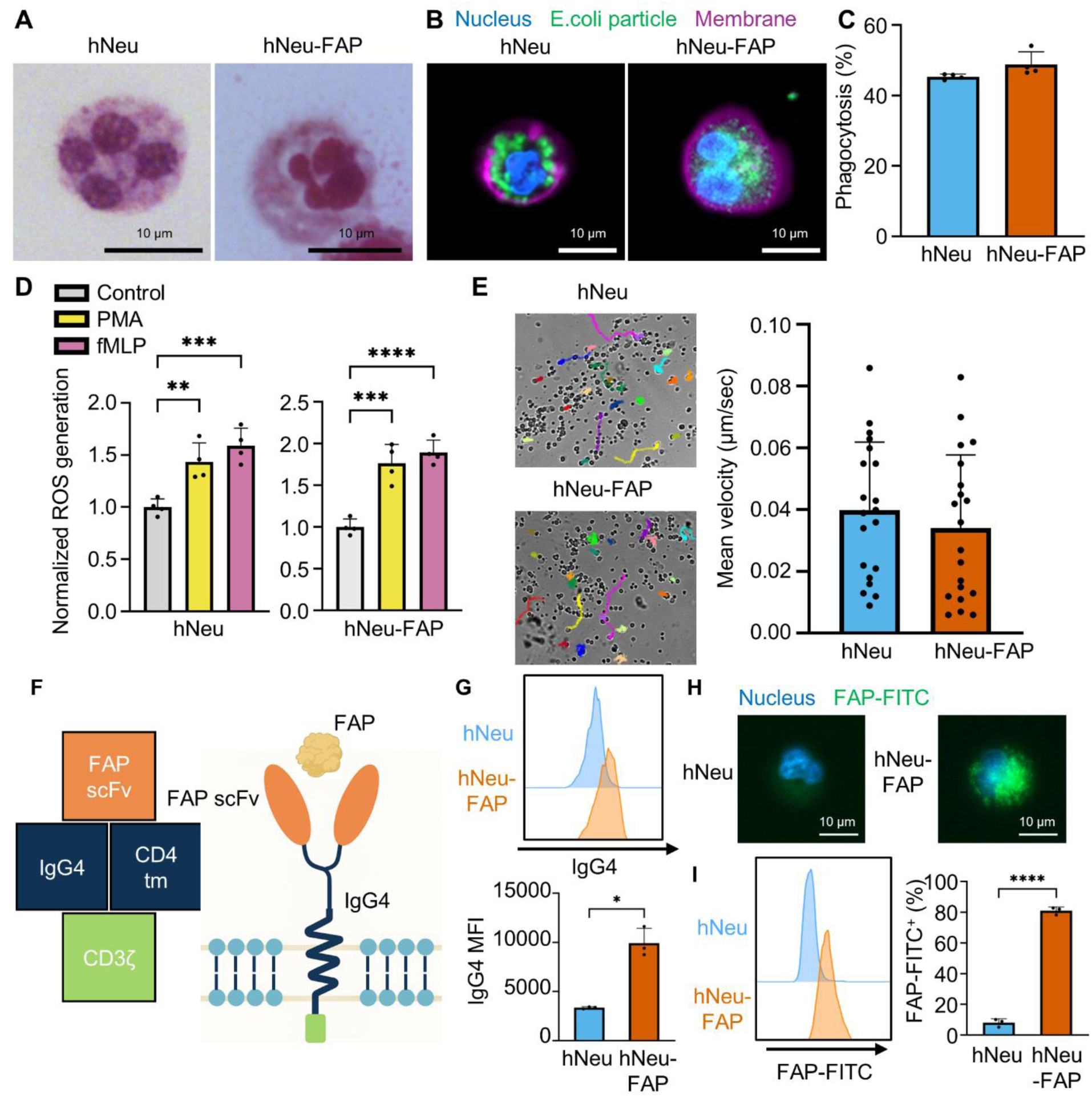
In vitro functional characterization of hNeu and hNeu-FAP. (A) Giemsa staining of hNeu and hNeu-FAP demonstrating characteristic multilobed nuclei. Scale bars, 10 µm. (B) Representative IF images of hNeu and hNeu-FAP following the phagocytosis of fluorescent E. coli bioparticles (3 h incubation). Scale bars, 10 µm. (C) Flow cytometry quantification of fluorescent E. coli bioparticle phagocytosis after 3 h of incubation. (D) Reactive oxygen species (ROS) generation in hNeu and hNeu-FAP following a 30-min stimulation with 1 µM PMA or fMLP, measured via CellROX Green. (E) Mean migration velocity quantified from 90-min time-lapse imaging. (F) Schematic representation of the FAP CAR construct, comprising an anti-FAP scFv, IgG4 hinge, CD4 transmembrane domain, and CD3ζ signaling domain. (G) Flow cytometry histogram and median fluorescence intensity (MFI) demonstrating increased IgG4 hinge expression in hNeu-FAP. (H) Representative fluorescence microscopy images and (I) flow cytometry quantification of FAP-FITC binding/uptake in hNeu-FAPs following a 24-h incubation. Scale bars, 10 µm. Data are represented as mean ± SD. Statistical significance was determined using unpaired Student’s t-tests for (C), (G), and (I) (n = 3 biological replicates) and for (E) (n = 20 cells per group), and one-way ANOVA with Tukey’s post hoc analysis for (D) (n = 4 biological replicates). *P < 0.05, **P < 0.01, ***P < 0.001, ****P < 0.0001.

We next evaluated innate effector functions to assess whether particle uptake, stimulus-induced ROS production, and motility were retained. Phagocytic capacity was tested by incubating cells with pHrodo-conjugated E. coli BioParticles for 3 hours; confocal microscopy and quantitative flow cytometry showed comparable uptake between hNeu and hNeu-FAP (**Figure 2B, C**), supporting retention of non-specific particle uptake under the assay conditions. Intracellular reactive oxygen species (ROS) production—a hallmark of respiratory burst activation—was evaluated using CellROX Green following 30-minute stimulation with phorbol 12-myristate 13-acetate (PMA) or N-formylmethionyl-leucyl-phenylalanine (fMLP). Both hNeu and hNeu-FAP showed statistically significant increases in ROS generation relative to unstimulated controls (**Figure 2D**). Single-cell tracking during 90-minute time-lapse microscopy showed no significant difference in mean migration velocity between the two cohorts under the under-agarose assay conditions containing fMLP (**Figure 2E**).

Bulk RNA sequencing showed high concordance in global gene-expression patterns between hNeu and hNeu-CAR (r = 0.9638, R^2^ = 0.9288; **Supplementary Figure 3A**). Analysis of canonical granulocytic transcripts confirmed consistent and abundant expression of myeloid lineage-associated genes (SPI1, CSF3R), granule enzymes (MPO, LYZ), and surface-marker-associated transcripts (ITGAM [CD11b], FUT4 [CD15], FCGR3A/B [CD16], CEACAM8 [CD66b], and CXCR2) across independent biological replicates (**Supplementary Figure 3B**).

We next assessed the surface display and antigen-binding fidelity of the integrated FAP CAR construct, which comprises an anti-FAP scFv, an IgG4 hinge/spacer, a CD4 transmembrane domain, and a CD3ζ intracellular activation domain (**Figure 2F**). Flow cytometric detection of the extracellular IgG4 hinge confirmed prominent surface CAR expression on hNeu-FAP compared to unmodified hNeu (**Figure 2G**). Following incubation with FAP-FITC, hNeu-FAP showed increased cell-associated fluorescence by microscopy (**Figure 2H**) and flow cytometry (**Figure 2I**), consistent with FAP binding and/or uptake. These results indicate that hNeu-FAP expressed the FAP CAR and retained the neutrophil functions examined here.

### 3.4 Polarization-Associated Changes in CAR-Expressing Neutrophils

We next examined differentially expressed genes in the FAP-CLTX dual CAR neutrophils (hNeu-CAR) RNA-seq dataset, focusing on pathways potentially relevant to the inflammatory and fibrotic cardiac environment [15]. Evaluation of the top differentially expressed genes (DEGs) via a global RNA-seq heatmap (**Figure 3A**) identified changes in inflammatory, granule-associated, and matrix-remodeling transcripts. Engineered hNeu-CAR cells exhibited upregulated expression of genes associated with extracellular matrix (ECM) remodeling, such as ADAMTS5, a metalloproteinase capable of cleaving fibrotic proteoglycans like versican and aggrecan. hNeu-CAR also showed increased expression of CNR2 (Cannabinoid Receptor 2), a candidate regulator of inflammatory responses, together with the macrophage-associated transcript ADGRE1. Concurrently, the engineered cells showed decreased expression of granule-associated components (CAMP, DEFA4) and the pro-fibrotic TGF-beta target CILP, suggesting altered granule-associated and matrix-related expression that could be relevant to fibroblast interactions. To further examine these transcriptional differences, we analyzed selected functional gene groups across the two biological replicates (**Figure 3B**). The hNeu-CAR cells showed higher expression of IRAK4, P2RY14, and HIF3A within the ‘Danger Sensing’ group. The ‘Tissue Infiltration’ group also showed higher expression of CXCL12, DOCK11, and ADAM8, which were grouped by putative roles in migration and matrix interactions. In contrast, the engineered cells exhibited reduced expression in the ‘N2-Associated’ block, with lower endogenous levels of ARG1, MMP8, CHI3L1, GPNMB, and the cardiac fibrosis-associated factor SERPINE1 (PAI-1) compared to unengineered controls. Volcano and MA plots summarized the magnitude and statistical significance of these transcriptional differences (**Figure 3C, D**). The MA plot demonstrated that the downregulated remodeling-associated targets (ARG1, MMP8, SERPINE1) are relatively abundant transcripts in the baseline unengineered hPSC-neutrophils. Lower OLFM4 and TUBB1 expression also suggested differences in differentiation-associated features. With respect to mediators relevant to inflammation and vascular remodeling, IL1B and VEGFA were also lower in the CAR cohort, providing candidates for future protein-level and functional analyses. IL12B remained largely unchanged and positioned in the low-expression region on the MA plot, indicating that the selected N1-associated genes did not uniformly increase. CXCL12, IRAK4, and TASL were also relatively abundant transcripts with higher expression in hNeu-CAR. We also examined selected N1/N2-associated transcripts in the RNA-seq data (**Figure 3E**). Flow cytometry further showed higher iNOS and lower Arginase-1 expression in hNeu-FAP than in hNeu (**Figure 3F**). The direction of these protein-level changes is consistent with the N1-associated features examined in our previous CAR-neutrophil work [21]. Together, these data show changes in selected polarization-associated features of CAR-expressing neutrophils. Whether these changes are maintained in the injured cardiac environment remains to be determined.

**Figure 3.**
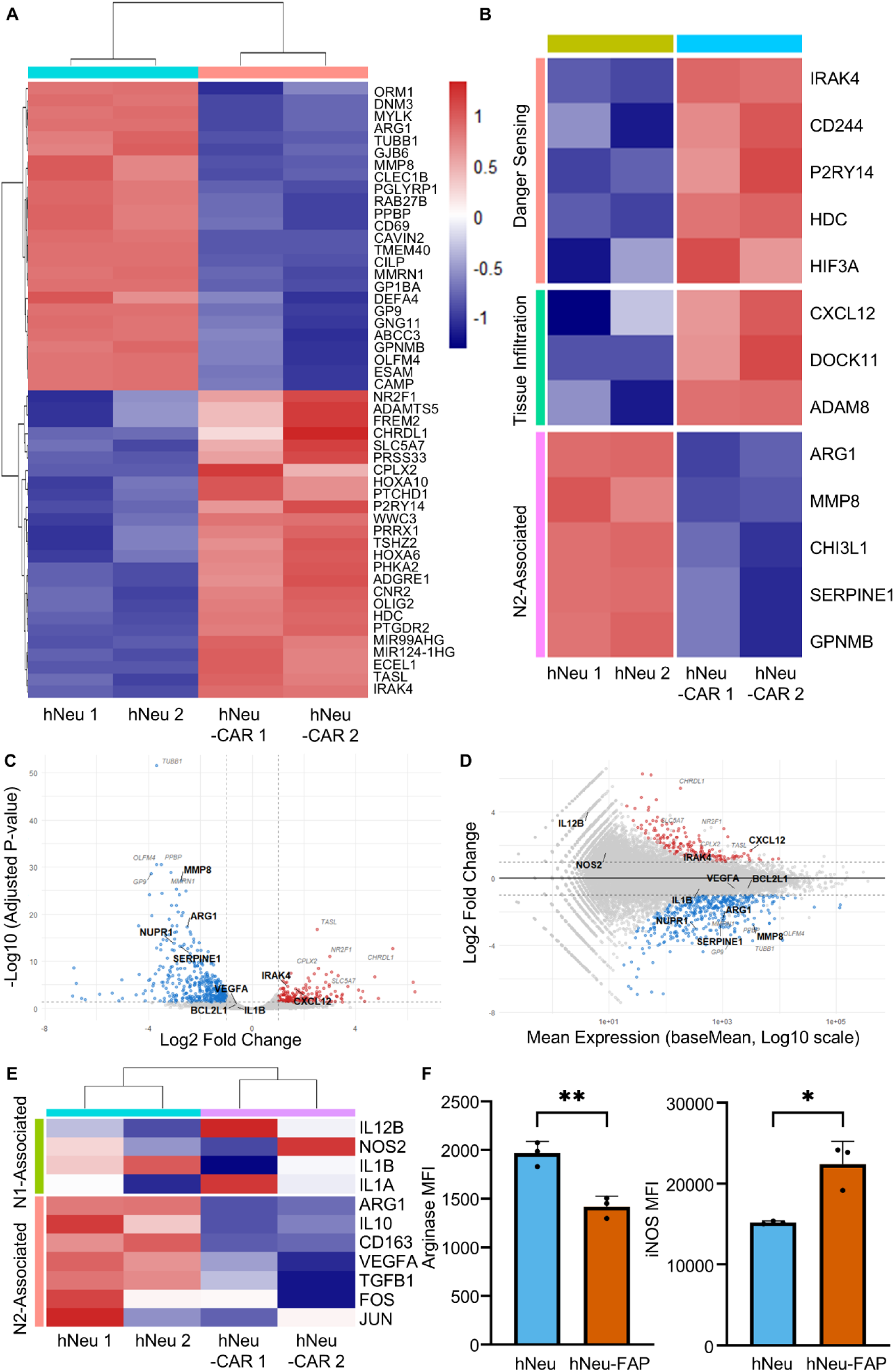
Polarization-associated features of CAR-expressing neutrophils. (A) Heatmap of the top 25 upregulated and 25 downregulated differentially expressed genes (DEGs) identified via bulk RNA-seq of hNeu and FAP-CLTX dual CAR neutrophils (hNeu-CAR). (B) Heatmap analysis of DEGs categorized by danger sensing, tissue infiltration, and N2/pro-fibrotic signaling. (C) Volcano plot displaying DEGs between hNeu and hNeu-CAR, highlighting significantly upregulated genes (red; including IRAK4 and CXCL12) and downregulated genes (blue; including ARG1, MMP8, and SERPINE1) based on adjusted P-value vs. Log2 Fold Change. Dashed lines denote significance and fold-change thresholds. (D) MA plot illustrating Log2 Fold Change against normalized mean expression (baseMean, Log10 scale), highlighting abundant downregulated pro-fibrotic transcripts (blue) alongside key upregulated effector/danger-sensing transcripts (red) and non-significant transcripts (grey). (E) Heatmap analysis of genes associated with N1 and N2 neutrophil polarization. (F) Flow cytometry quantification (MFI) of Arginase1 (N2 marker) and iNOS (N1 marker) in hNeu and hNeu-FAP. Data are represented as mean ± SD. Statistical significance was determined by unpaired Student’s t-tests comparing hNeu vs. hNeu-FAP (n = 3 biological replicates). *P < 0.05, **P < 0.01, ***P < 0.001, ****P < 0.0001.

### 3.5 Targeting of Activated Epicardial Fibroblasts in 2D and 3D Microenvironments

Epicardium-derived cells represent a major source of activated myofibroblasts driving pathological cardiac remodeling and scar deposition [30]. To model this target population in vitro, we differentiated epicardial fibroblasts from hPSCs [25] and activated them with TGF-beta and serum to generate activated fibroblasts characterized by FAP and VIMENTIN staining (**Supplementary Figure 4**).

To assess the cytotoxic activity of hNeu-FAP against activated cardiac fibroblasts, 2D co-culture assays were performed with GFP- or luciferase-expressing target cells. Fluorescence imaging revealed a reduction of GFP+ fibroblasts following overnight co-culture with hNeu-FAP (**Figure 4A**). Quantitative cell viability assays across various effector-to-target (E:T) ratios demonstrated dose-dependent cytolytic activity by hNeu-FAP, reducing fibroblast viability to approximately 42% at a 10:1 E:T ratio, whereas unmodified hNeu exhibited negligible baseline cytotoxicity (**Figure 4B**). Time-lapse microscopy showed hNeu-FAP contacting and disrupting target fibroblasts (**Supplementary Video 1, 2**).

**Figure 4.**
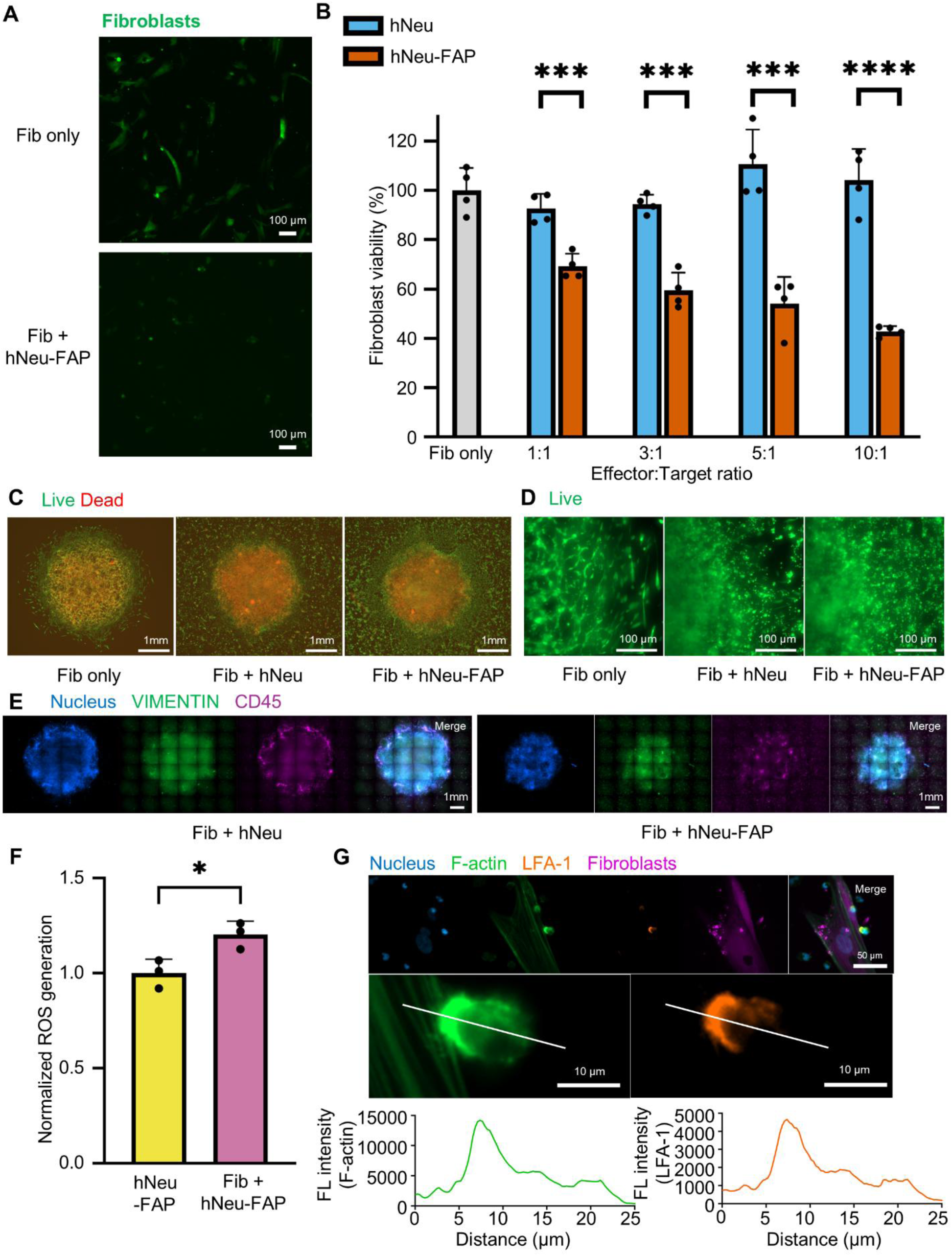
hNeu-FAP targets activated epicardial fibroblasts in 2D and 3D in vitro systems. (A) Representative fluorescence images of GFP-expressing activated epicardial fibroblasts co-cultured with or without hNeu-FAP. Scale bars, 100 µm. (B) Normalized viability of activated fibroblasts following co-culture with hNeu or hNeu-FAP at the indicated effector-to-target (E:T) ratios. (C) Fluorescence images (Calcein AM/PI staining) of 3D epicardial fibroblast structures co-cultured with hNeu or hNeu-FAP. Scale bars, 1 mm. (D) Representative fluorescence images of live cells at the periphery of the 3D fibroblast structures. Scale bars, 100 µm. (E) Maximum intensity projections of 3D epicardial fibroblasts co-cultured with hNeu or hNeu-FAP, co-stained for fibroblasts (VIMENTIN) and neutrophils (CD45). (F) Normalized CellROX fluorescence in hNeu-FAP alone and in fibroblast–hNeu-FAP co-cultures after 4 h. (G) Confocal imaging revealing polarized F-actin and LFA-1 accumulation at the hNeu-FAP–fibroblast interface, possibly indicating immune-synapse-like structural formations. Fibroblasts were stained with CellTrace-APC immediately prior to co-culture with hNeu-FAP. Scale bars, 50 or 10 µm. Data are represented as mean ± SD. Statistical significance was determined by unpaired Student’s t-tests for the indicated comparisons (n = 3 or 4 biological replicates). *P < 0.05, **P < 0.01, ***P < 0.001, ****P < 0.0001.

We next evaluated targeting efficacy within a more complex tissue architecture by generating 3D fibroblast micro-tissues. Live/dead viability staining (Calcein AM/PI) showed that control micro-tissues maintained dense, interconnected networks of elongated spindle-shaped fibroblasts (**Figure 4C, D**). In contrast, co-culture with hNeu-FAP caused disruption of the cellular network and loss of cell viability, particularly along the outer perimeter of the 3D structures (**Figure 4C, D**). Whole-mount confocal tile scanning for VIMENTIN (fibroblasts) and CD45 (neutrophils) showed CD45+ hNeu-FAP signal over central regions of the projected micro-tissue images, accompanied by structural compaction and reduced overall spheroid dimensions relative to unmodified hNeu co-cultures (**Figure 4E**). Because ROS generation has been implicated in hPSC-neutrophil-mediated target-cell interactions [21], we next measured CellROX fluorescence during fibroblast co-culture. Co-incubation of hNeu-FAP with target fibroblasts was associated with significantly higher CellROX fluorescence than hNeu-FAP monocultures (**Figure 4F**). Confocal imaging and fluorescence intensity line scans showed enrichment of F-actin and LFA-1 at hNeu-FAP–fibroblast contact sites, consistent with synapse-like organization (**Figure 4G**). Together, these findings demonstrate fibroblast-directed activity of hNeu-FAP in both 2D and 3D environments, accompanied by increased CellROX fluorescence and synapse-like organization at effector-target contact sites.

### 3.6 hNeu-FAP Cells Target Activated Fibroblasts in 3D Cardiac Organoids

To evaluate potential bystander effects and examine therapeutic performance within a physiologically relevant tissue architecture, we engineered multi-lineage 3D human cardiac-fibroblast organoids (hCFOs) (**Figure 5A**). These microtissues were formed by co-aggregating hPSC-derived cardiomyocytes (CMs) with GFP-expressing, activated epicardial-derived fibroblasts, followed by co-culture with either unmodified hNeu or hNeu-FAP (**Figure 5A**). Live-cell fluorescence imaging showed significantly lower GFP signals in hCFOs co-cultured with either hNeu or hNeu-FAP than in untreated controls (**Figure 5B, C**).

**Figure 5.**
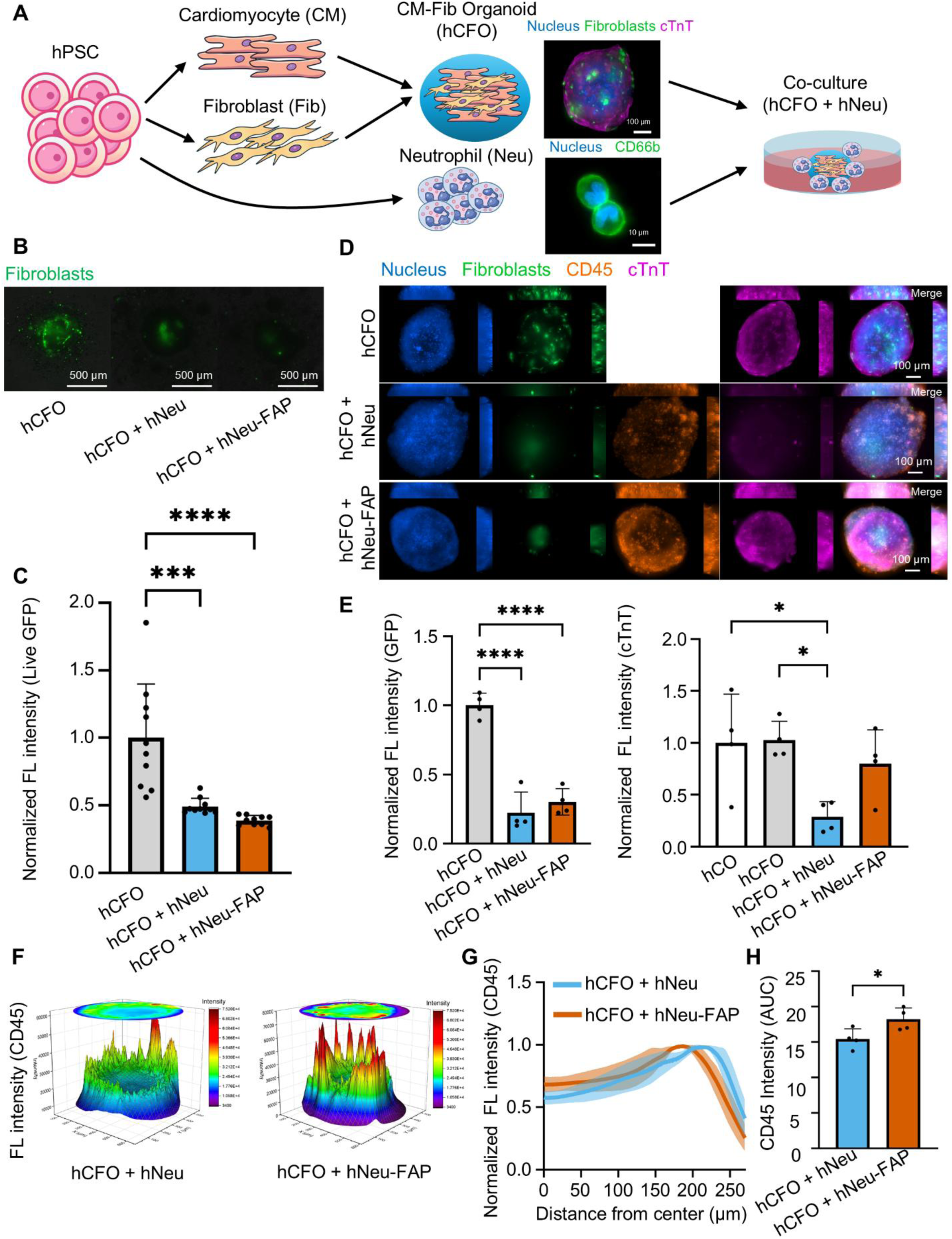
hNeu-FAP targets activated epicardial fibroblasts within a 3D cardiac organoid model. (A) Schematic overview of the differentiation of cardiomyocytes (CMs), epicardial fibroblasts (Fibs), and neutrophils (hNeu) from hPSCs, the subsequent formation of CM-Fib organoids (hCFOs), and representative fluorescence images of hCFOs and hNeu. (B) Representative live-cell images of hCFOs containing GFP-expressing epicardial fibroblasts co-cultured with hNeu or hNeu-FAP. (C) Quantification of live GFP expression. (D) Representative maximum intensity projections of hCFOs co-cultured with hNeu or hNeu-FAP (Green: fibroblasts; Orange: CD45/hNeu; Magenta: cTnT/CMs). (E) Quantification of GFP (fibroblasts) and cTnT (CMs) fluorescence intensity in hCFOs. (F) Fluorescence-intensity surface plot of CD45 derived from sum-intensity projections of confocal z-stacks of hCFOs co-cultured with hNeu or hNeu-FAP. (G) Normalized fluorescence intensity radial profiles of the CD45 sum-intensity projections. (H) Quantification of CD45 intensity near the projected organoid center (within 10% of the radius), calculated by the area under the curve (AUC) of the radial profiles. Data are represented as mean ± SD. Statistical significance was determined by unpaired Student’s t-tests comparing hNeu vs. hNeu-FAP for (H) (n = 4 organoids per group) and by one-way ANOVA with Tukey’s post hoc analysis for (C) (n = 10 organoids per group) and (E) (n = 4 organoids per group). *P < 0.05, **P < 0.01, ***P < 0.001, ****P < 0.0001.

To examine whether the reduction in fibroblast-associated signal was accompanied by changes in adjacent cardiac parenchyma, organoids were fixed and immunostained for cardiac troponin T (cTnT; cardiomyocytes), CD45 (neutrophils), and GFP (fibroblasts) (**Figure 5D**). Quantitative confocal analysis confirmed a significant reduction of GFP fluorescence in both neutrophil-treated cohorts (**Figure 5E, left**). However, analysis of cTnT expression revealed a different pattern across the treatment conditions (**Figure 5E, right**). While co-culture with unmodified hNeu led to a statistically significant decrease in cTnT intensity compared with untreated organoid controls (P < 0.05), hCFOs treated with hNeu-FAP showed no significant change in cTnT intensity (Figure 5E, right). These results suggest differential effects of unmodified and FAP CAR neutrophils on the cardiomyocyte-associated signal within the organoids.

We next evaluated whether CAR targeting influenced the radial distribution of neutrophil-associated fluorescence within the dense 3D organoid microenvironment (**Figure 5F–H**). Surface plots generated from sum-intensity projections of confocal z-stacks displayed localized CD45 fluorescence intensity with higher peaks across the organoid area in the hNeu-FAP group compared to unmodified hNeu (**Figure 5F**). To compare the radial signal patterns after accounting for differences in peak intensity, radial fluorescence intensity profiles were normalized to each condition’s peak peripheral accumulation (**Figure 5G**). While both groups displayed maximal CD45 accumulation at the outer rim (approximately 175–225 µm from the center), hNeu-FAP maintained a higher relative signal toward the projected center (0–150 µm) (**Figure 5G**). Quantitative integration of these normalized radial distributions via area under the curve (AUC) showed a significant difference in the central CD45 fluorescence measure for hNeu-FAP compared to unmodified hNeu (P < 0.05; **Figure 5H**). Together, these observations suggest a difference in the radial distribution of CD45 fluorescence within the projected microtissue area, while depth-resolved imaging would be needed to determine whether this reflects greater penetration into the organoid interior.

Finally, to determine whether the effects observed with differentiated neutrophils were also present at an earlier developmental stage, we performed parallel co-culture experiments using Day 12 hematopoietic progenitor cells (hHPCs and hHPC-FAP) (**Supplementary Figure 5**). Live-cell imaging revealed only a marginal reduction in live GFP signal across both progenitor-treated groups (**Supplementary Figure 5A, B**). Consistent with these modest live-cell changes, fixed immunostaining and confocal quantification demonstrated no significant alterations in either GFP (fibroblast) or cTnT (cardiomyocyte) intensity compared to untreated controls (**Supplementary Figure 5C, D**). These findings suggest that the effects observed with differentiated neutrophils were not reproduced at the progenitor stage under the conditions tested.

### 3.7 Single-Cell and Single-Nucleus Transcriptomics Map Immune-Associated Expression and Fibroblast Activation in Murine Cardiac Injury

For FAP-directed neutrophil therapy, treatment timing must account for both the arrival of effector cells in injured myocardium and the availability of target antigens. We therefore analyzed public single-cell and single-nucleus RNA-sequencing datasets across multiple cardiac injury models to compare temporal patterns of immune-associated expression and fibroblast activation markers and inform the choice of an intervention time point (**Figure 6 and Supplementary Figure 6**).

**Figure 6.**
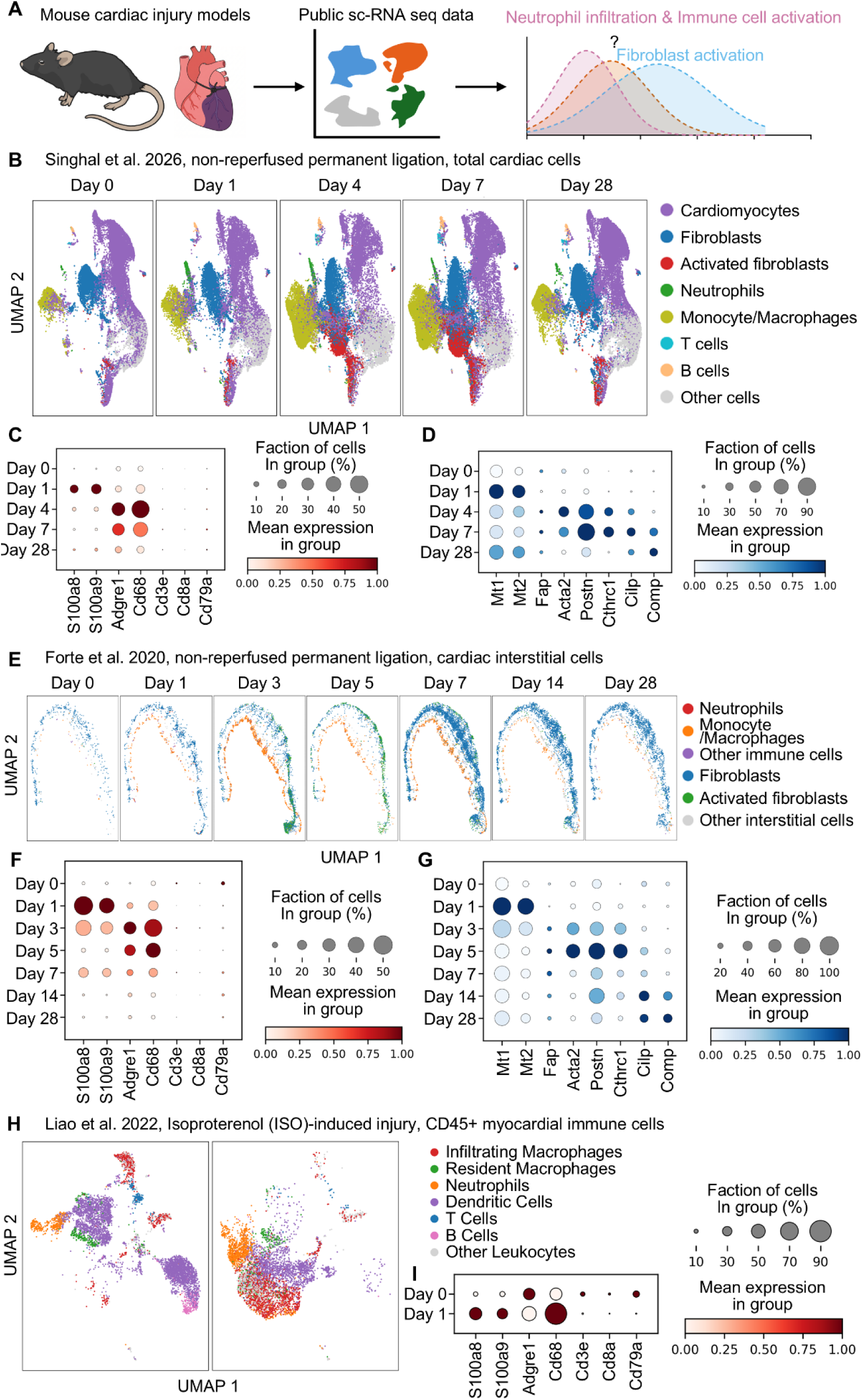
Single-cell and single-nucleus RNA-seq reveals temporal immune- and fibroblast-associated expression patterns in murine cardiac injury models. (A) Schematic representation of the computational workflow for evaluating immune recruitment and stromal activation across public cardiac single-cell datasets. (B) UMAP embedding of sampled cardiac cells or nuclei across five time points (Day 0, 1, 4, 7, and 28) following permanent coronary artery ligation (Singhal et al., 2026), annotated via automated module scoring into parenchymal, stromal, and immune subsets. (C) Gene-wise scaled dot plot showing temporal expression of markers associated with canonical immune lineages, including neutrophils (S100a8, S100a9), monocytes/macrophages (Adgre1, Cd68), T cells (Cd3e, Cd8a), and B cells (Cd79a). (D) Dot plot showing fibroblast-associated expression patterns, including early cellular stress response (Mt1, Mt2), active extracellular matrix remodeling (Fap, Acta2, Postn, Cthrc1), and late matrifibrocyte maturation (Cilp, Comp). (E) UMAP projection of cardiac interstitial cells across seven post-injury stages (Days 0 to 28) in a non-reperfused permanent ligation model (Forte et al., 2020), displaying marker-based annotations of fibroblast and immune compartments. (F) Gene-wise scaled dot plot depicting temporal granulocyte- and monocyte-associated expression and (G) corresponding fibroblast activation-associated expression in the Forte et al. dataset. (H) UMAP embedding of sorted CD45+ myocardial leukocytes isolated from an isoproterenol (ISO)-induced injury model (Liao et al., 2022), displaying marker-based myeloid and lymphoid annotations including putative infiltrating (Ccr2, Ly6c2) and resident (Cx3cr1, Mrc1) macrophages alongside neutrophil-associated cells. (I) Dot plot illustrating the rapid shift in immune signature expression between uninjured baseline (Day 0) and acute stress (Day 1). Circle size indicates the fraction of sampled cells or nuclei expressing each gene within the plotted group; color denotes the mean expression rescaled from 0 to 1 separately for each gene across groups. Dot plots summarize the sampled population rather than a fibroblast-only subset and do not directly measure absolute cell recruitment.

We first mapped cellular dynamics in a permanent coronary ligation myocardial infarction (MI) model across an acute-to-chronic timeline (Days 0, 1, 4, 7, and 28; Singhal et al., 2026 [31]) **(Figure 6B–D**). UMAP visualization and marker profiles across sampled cardiac cells and nuclei showed prominent neutrophil-associated expression (S100a8, S100a9) at Day 1 post-injury, followed by stronger monocyte/macrophage-associated signatures (Adgre1, Cd68) at Days 4 and 7 (**Figure 6B, C**). Alongside baseline fibroblast markers (Dcn, Pdgfra), the expression panels showed early stress-associated metallothionein expression (Mt1, Mt2) at Day 1 (**Figure 6D**). This early expression pattern was followed by prominent fibroblast activation-associated expression of Fap, Acta2, Postn, and Cthrc1 between Day 4 and Day 7 (**Figure 6D**). By Day 28, the panels showed later matrifibrocyte-associated Cilp and Comp expression (**Figure 6D**). A similar ordering of expression patterns was observed in an independent cardiac interstitial dataset following non-reperfused permanent ligation (Forte et al., 2020 [32]) (**Figure 6E–G**). Neutrophil-associated S100a8/S100a9 expression was prominent at Day 1, preceding stronger monocyte/macrophage-associated expression at Days 3 to 5 (**Figure 6E, F**). In the same dataset, fibroblast activation markers (Fap, Acta2, Postn, Cthrc1) were prominent at Day 5, whereas matrifibrocyte-associated Cilp and Comp expression was evident at Days 14 and 28 (**Figure 6G**). Similarly, in a non-ischemic, chemical stress model induced by isoproterenol (ISO; Liao et al., 2022 [33]), profiling of CD45+ leukocytes showed increased neutrophil-associated (S100a8, S100a9) and macrophage-associated (Cd68, Adgre1) expression within 24 hours post-administration (**Figure 6H, I**).

To evaluate whether these early immune-associated and subsequent remodeling-associated patterns extend across injury contexts, we examined additional acute and chronic cardiac injury models (**Supplementary Figure 6**). In an ischemia-reperfusion (I/R) model (Lin et al., 2023 [34]), Day 1 was characterized by prominent neutrophil-associated expression accompanied by stress-associated expression (Mt1, Mt2) and early emergence of Fap expression (**Supplementary Figure 6A**). Under chronic pressure overload and neurohormonal stress—including angiotensin II/phenylephrine infusion (AngII/PE; Boccella et al., 2025 [35]) and transverse aortic constriction (TAC; Alexanian et al., 2021 [36])—hearts displayed persistent immune-associated expression accompanied by fibroblast activation-associated programs marked by Fap, Postn, and Meox1 (**Supplementary Figure 6B, C**). Collectively, these temporal expression profiles are consistent with an early neutrophil-associated response after acute injury (Day 1), followed by increased expression of markers associated with FAP-positive fibroblast activation over Days 3–7. Because the dot plots summarize sampled populations, these patterns may reflect both changes in cell composition and changes in expression within cells. These temporal patterns informed the choice of Day 4 for administration, when fibroblast activation-associated expression was increasing.

### 3.8 In Vivo Myocardial Localization of Endogenous and Adoptively Transferred Neutrophils

To examine the early neutrophil response suggested by the transcriptomic analyses, we assessed neutrophil-associated signals in mouse hearts 24 h after ISO-induced injury (**Figure 7**). Immunofluorescence analysis of heart sections harvested 24 h post-ISO challenge showed greater Ly6G staining in ISO-treated hearts than in PBS-treated controls (**Figure 7A**). Because non-specific vascular pooling can confound tissue-infiltrating leukocyte quantification, extensive transcardial perfusion was performed prior to tissue collection to flush out residual intravascular cells. Flow cytometric analysis of CD45+-enriched cardiac leukocytes showed a numerically higher frequency of Ly6G+ neutrophils in ISO-injured hearts than in vehicle-treated controls, although the difference did not reach statistical significance (P = 0.0962; **Figure 7B**).

**Figure 7.**
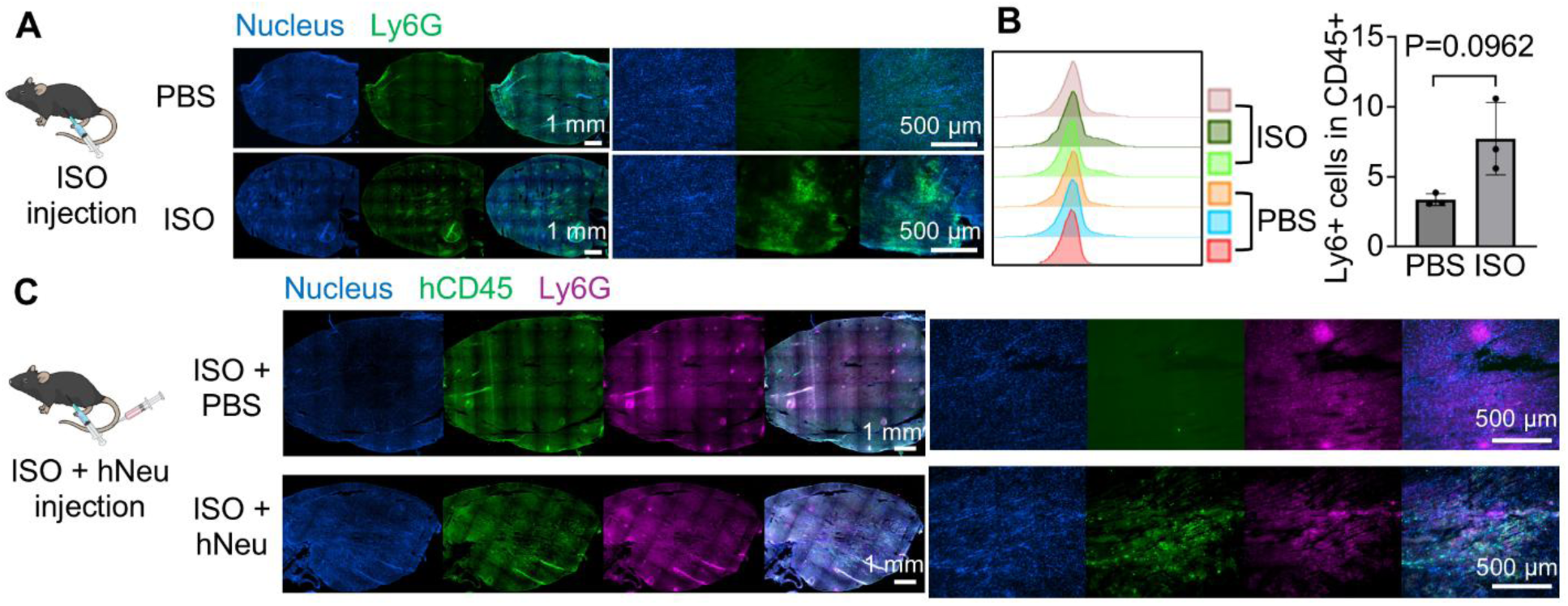
Myocardial localization of endogenous and adoptively transferred neutrophils following acute cardiac injury. (A) Representative IF images of Ly6G+ murine neutrophils infiltrating the heart 24 h post-ISO injection. Scale bars, 1 mm and 500 µm. (B) Flow cytometry analysis showing the percentage of Ly6G+ cells in CD45+ cells from PBS- and ISO-treated mouse hearts (n = 3 mice per group; P = 0.0962). (C) Representative IF images showing Ly6G+ murine neutrophils and human CD45+ hNeu in myocardial sections of immunosuppressed mice 24 h following ISO and hNeu administration. Scale bars, 1 mm and 500 µm.

We next asked whether systemically administered hNeu could also be detected in the injured myocardium (**Figure 7C**). Mice were temporarily immunosuppressed with cyclosporine A (CsA) and intravenously injected with 8 × 10^6 hNeu concurrent with ISO administration (**Figure 7C, left**). At 24 h post-injection, animals were transcardially perfused to clear circulating cells, and excised myocardium was co-stained for human CD45 (hCD45) and murine Ly6G (**Figure 7C**). Immunofluorescence imaging of whole-heart cross sections demonstrated hCD45+ signal within myocardial sections of hNeu-treated mice, whereas hearts from vehicle-injected animals (ISO + PBS) displayed negligible human CD45 background signal **(Figure 7C**). High-magnification regional scans showed that human CD45-positive signal was present in regions around endogenous Ly6G+ murine neutrophils (**Figure 7C, right**). Together, these results support the presence of systemically delivered hPSC-derived neutrophils in perfused ISO-injured hearts within 24 h, providing a rationale for their evaluation in cardiac injury without directly establishing the mechanism or specificity of myocardial homing.

### 3.9 In Vivo Evaluation of Cardiac Function and Fibrosis Following FAP CAR Neutrophil Administration Post-Myocardial Infarction

To evaluate the effects of FAP CAR neutrophils in vivo, we employed a permanent coronary artery ligation model of myocardial infarction (MI) in mice (**Figure 8A**). Based on the transcriptomic profiles showing increased fibroblast activation-associated expression between Days 3 and 7 after injury, we administered PBS, hNeu, or hNeu-FAP intravenously at Day 4 post-MI (**Figure 8A**). Cardiac performance was assessed by echocardiography prior to surgery (baseline) and at Day 7, with hearts collected at Day 14 for histological analysis (**Figure 8A**).

**Figure 8.**
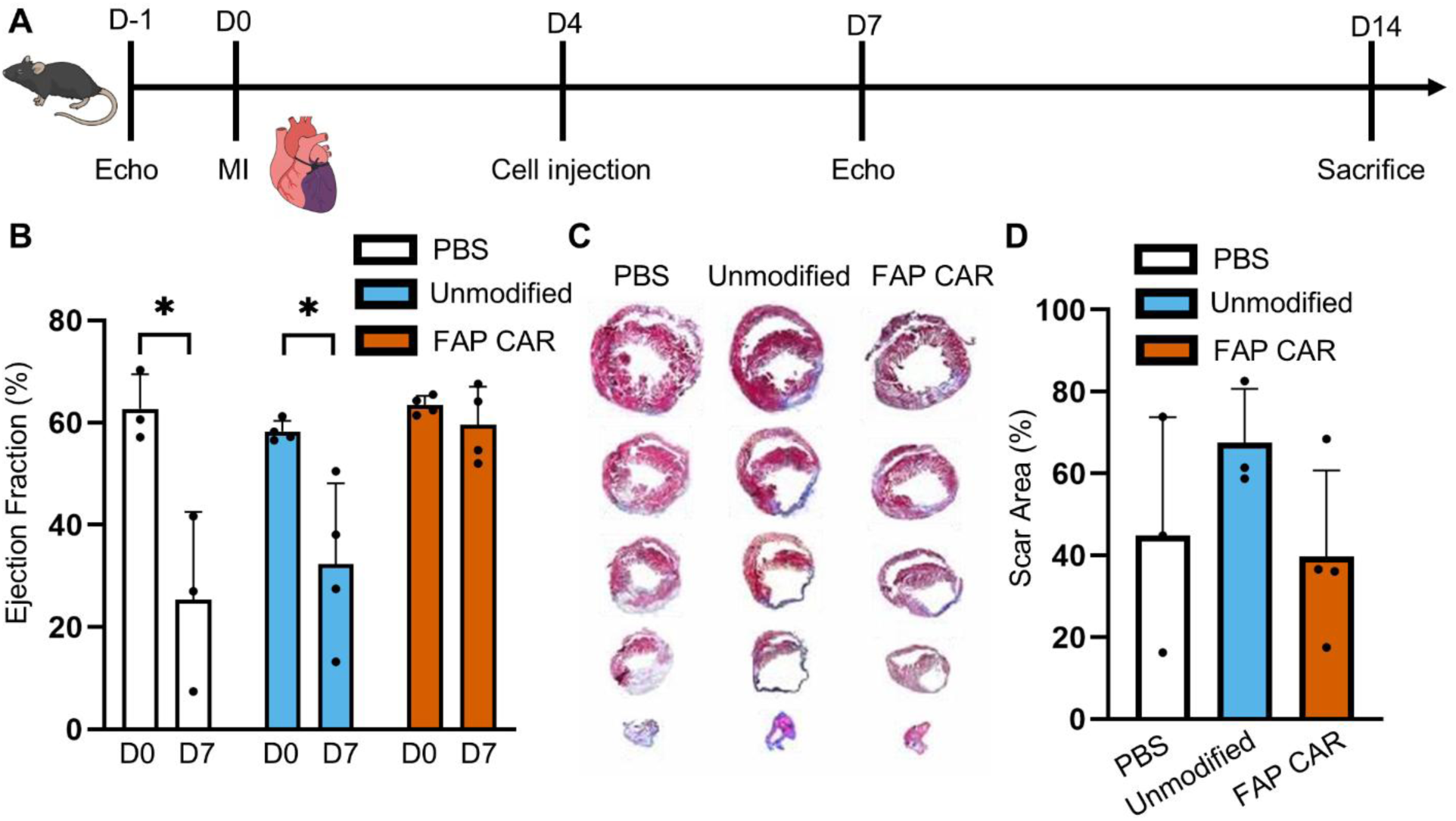
In vivo evaluation of hNeu-FAP in a murine myocardial infarction (MI) model. (A) Experimental timeline detailing MI induction (permanent ligation), cell injection (1 × 10^7 cells/mouse), and echocardiography (echo) analysis. (B) Ejection fraction of mice treated with PBS (control), hNeu, or hNeu-FAP at day 0 and day 7 post-MI induction. (C) Histological analysis of sectioned hearts collected at day 14 post-MI. (D) Quantification of the scar area in sectioned hearts at day 14 post-MI. Data are represented as mean ± SD.

Serial echocardiographic assessment showed a decline in left ventricular ejection fraction (LVEF) by Day 7 post-MI in both control cohorts, with animals receiving vehicle (PBS) or unmodified neutrophils exhibiting substantial functional loss from baseline (PBS: 62.7% ± 6.8% at D0 vs. 25.4% ± 17.2% at D7, P < 0.05; hNeu: 58.3% ± 2.1% at D0 vs. 32.3% ± 15.8% at D7, P < 0.05; **Figure 8B**). In the hNeu-FAP group, in contrast, mean LVEF was 63.4% ± 1.8% at baseline and 59.6% ± 7.4% at Day 7 (**Figure 8B**).

To determine the impact of treatment on cardiac remodeling and fibrosis, hearts were collected at Day 14 post-MI and subjected to Masson’s trichrome staining across serial transverse cross-sections (**Figure 8C**). Myocardial sections from PBS- and unmodified neutrophil-treated mice exhibited anterior wall thinning, chamber dilatation, and collagenous scarring (**Figure 8C**). Hearts treated with FAP CAR neutrophils showed thicker left ventricular walls and less collagen staining in the infarct region (**Figure 8C**). Morphometric quantification showed a reduction in overall fibrotic scar area in mice treated with FAP CAR neutrophils compared to those receiving unmodified neutrophils or vehicle control (**Figure 8D**).

## 4. Discussion

Cardiac fibrosis remains a major driver of adverse ventricular remodeling and progression to heart failure, yet current clinical therapies primarily modulate upstream neurohormonal signaling rather than directly eliminating pathogenic myofibroblasts or removing established ECM accumulation [1,2]. While CAR T cells targeting FAP have demonstrated the feasibility of ablating fibrogenic stromal cells [5,6], translating lymphoid cell therapies to chronic non-malignant conditions is hindered by autologous manufacturing bottlenecks, long-term lymphoid persistence, and the potential for on-target injury to FAP-expressing healthy stromal niches [7,8,37]. CAR-macrophages and CAR-monocytes provide tissue-infiltrating and phagocytic effector functions [10–12], although their phenotypic plasticity allows local myocardial signals to influence their inflammatory and matrix-remodeling activities [14–16].

Here, we developed an hPSC-derived FAP CAR neutrophil platform that incorporates progenitor banking. We first examined whether chimeric receptor expression altered basic neutrophil functions. Differentiated hNeu-FAP retained characteristic polymorphonuclear morphology, motility under fMLP-containing assay conditions, and preserved respiratory burst capacity following chemical stimulation. These results indicate that CAR expression did not measurably impair the neutrophil functions examined here. Staining against the extracellular human IgG4 spacer showed CAR expression on differentiated hNeu-FAP cells. Furthermore, upon incubation with recombinant FAP-FITC, hNeu-FAP showed significantly greater cell-associated antigen fluorescence compared to unmodified neutrophils. Because professional phagocytes enhance endocytic uptake and target ingestion following CAR engagement [38,39], the increased FAP-FITC signal is compatible with receptor binding and/or internalization, although this soluble-antigen assay does not itself establish target-cell phagocytosis. Thus, FAP CAR integration endows hPSC-derived neutrophils with FAP-associated antigen recognition and activity against activated fibroblasts while retaining the granulocytic functions tested here.

Neutrophil-based therapies must also account for the potential consequences of dysregulated granulocyte activation. In chronic inflammatory conditions and myocardial infarction, uncontrolled neutrophil activation—marked by systemic degranulation, diffuse ROS release, microvascular plugging, and the cytotoxic extrusion of neutrophil extracellular traps (NETs)—could drive substantial collateral tissue damage [23,40]. Moreover, non-specific or premature proteolytic degradation of the provisional myocardial ECM can critically compromise ventricular wall integrity, elevating the risk of mechanical cardiac rupture [41]. Our findings link fibroblast-directed activity to contact-associated structural polarization, while the organoid experiments provide an initial assessment of accompanying changes in neighboring cardiomyocytes. When hNeu-FAP contacted target fibroblasts, F-actin and LFA-1 accumulated at the cell–cell interface in a synapse-like pattern. In 3D hCFOs, both neutrophil groups reduced GFP fluorescence, whereas a significant decrease in cTnT intensity relative to untreated controls was detected only with unmodified hNeu. Together, these findings suggest differential effects of hNeu and hNeu-FAP within the cardiac co-culture model. Further studies using direct cardiomyocyte viability measurements and spatially resolved ROS analysis will help define the target specificity and mechanisms underlying these effects.

Beyond target-cell interactions, the activity of innate effector cells in fibrotic tissue may also depend on their polarization state. Paralleling the macrophage M1/M2 framework, N1- and N2-associated neutrophil states have been described in relation to anti-tumor and pro-tumor activity, although this classification should not be treated as a universal anti-fibrotic/pro-fibrotic division [42,43]. In tissue repair and wound healing, local cues influence neutrophil behavior, while TGF-β has been implicated in N2-like polarization in tumor-associated settings [18,42]. In chronically inflamed or repairing tissues, factors such as Arginase-1, VEGF, and matrix-remodeling regulators, including MMP8 and SERPINE1, may influence inflammatory resolution and fibroblast activity in a context-dependent manner [1,43–45]. In our study, transcriptomic profiling of hNeu-CAR and flow cytometry of hNeu-FAP identified differences in several of these polarization-associated features. The hNeu-CAR showed increased expression of inflammatory signaling-associated transcripts (IRAK4, P2RY14) and migration- or matrix-associated genes (CXCL12, ADAM8, ADAMTS5), while hNeu-FAP neutrophils showed higher iNOS and lower Arginase-1 fluorescence. Concurrently, the CAR cells showed lower expression of several remodeling-associated transcripts, including ARG1, MMP8, CHI3L1, and SERPINE1. These observations build on our previous characterization of N1-associated features in CAR-engineered hPSC-derived neutrophils [21]. Whether these features persist and influence function in the injured myocardium remains unresolved.

Treatment timing is likely to be important because fibroblasts contribute to early wound stabilization before later scar formation. Cross-referencing public single-cell transcriptomic datasets across diverse cardiovascular injury models showed early neutrophil-associated expression at Day 1 post-injury, accompanied by stress-associated genes (Mt1, Mt2) and followed by increased fibroblast activation-associated expression (Fap, Acta2, Postn, Cthrc1) between Days 4 and 7 [31–33].

Guided by this temporal pattern, hNeu-FAP was administered at Day 4 post-MI, when fibroblast activation-associated expression was increasing and before the later Cilp/Comp-associated profile became prominent. The treated group showed higher Day 7 LVEF than PBS controls in the exploratory comparison, while the difference from unmodified hNeu and the Day 14 scar-area differences did not reach statistical significance. The short physiological lifespan of granulocytes provides a rationale for limiting prolonged effector exposure, although it does not replace direct evaluation of persistence, biodistribution, and acute inflammatory toxicity [18,19]. Host cardiac macrophages may subsequently clear apoptotic transferred neutrophils through efferocytosis [46,47].

Several aspects of the platform remain to be optimized. The first-generation CD3ζ signaling domain supported cytotoxic activity, but future screening of innate-specific intracellular signaling motifs could help optimize myeloid effector activity and cytokine release [48,49]. Cell storage represents another key translational consideration. Cryopreservation of Day 12 hematopoietic progenitors established a precursor-banking workflow that supports subsequent neutrophil differentiation rather than direct storage of a mature-cell product. Using hypoimmunogenic hPSC lines could broaden the allogeneic applicability of this precursor-banking approach. Transient in vivo engineering using mRNA-loaded nanocarriers [50] could be explored separately as an alternative approach.

## 5. Conclusion

In summary, this study evaluated hPSC-derived FAP CAR neutrophils as a cellular approach to cardiac fibrosis. By combining FAP-directed receptor engineering with hPSC differentiation, the platform generated neutrophils that showed dose-dependent activity against activated epicardial fibroblasts in 2D culture and reduced fibroblast-associated fluorescence in 3D cardiac organoids, without a significant reduction in cardiomyocyte-associated cTnT staining. F-actin and LFA-1 accumulated at hNeu-FAP–fibroblast contacts in a synapse-like pattern, and co-culture was accompanied by increased ROS-associated fluorescence. Transcriptomic and protein analyses also identified differences in selected inflammatory and polarization-associated features. In vivo, administration at Day 4 after myocardial infarction was associated with higher Day 7 LVEF than PBS treatment in an exploratory comparison, although evidence for a reduction in scar area remained inconclusive. The intermediate progenitor cryopreservation workflow also provides a route for cell banking before neutrophil differentiation, and the short lifespan of mature neutrophils warrants further study of treatment duration and safety. These findings support further studies of target dependence, cardiac remodeling, and therapeutic safety before extending the approach to other fibroproliferative diseases.

## 6. Acknowledgement

This work was supported by the National Institutes of Health (R37CA265926 to X.B., R35HL166557 to Y.N.H., and R35GM119787 to Q.D.), the National Science Foundation CAREER Award (CBET-2143064 to X.B.), and the American Heart Association (Postdoctoral Fellowship 23POST1025924 to G.J.; Postdoctoral Fellowship 25POST1373848 and Career Development Award 26CDA1609618 to S.F.).

