## Supplementary information for "Targeting Cardiac Fibrosis with Chimeric Antigen Receptor Neutrophils Derived from Human Pluripotent Stem Cells"

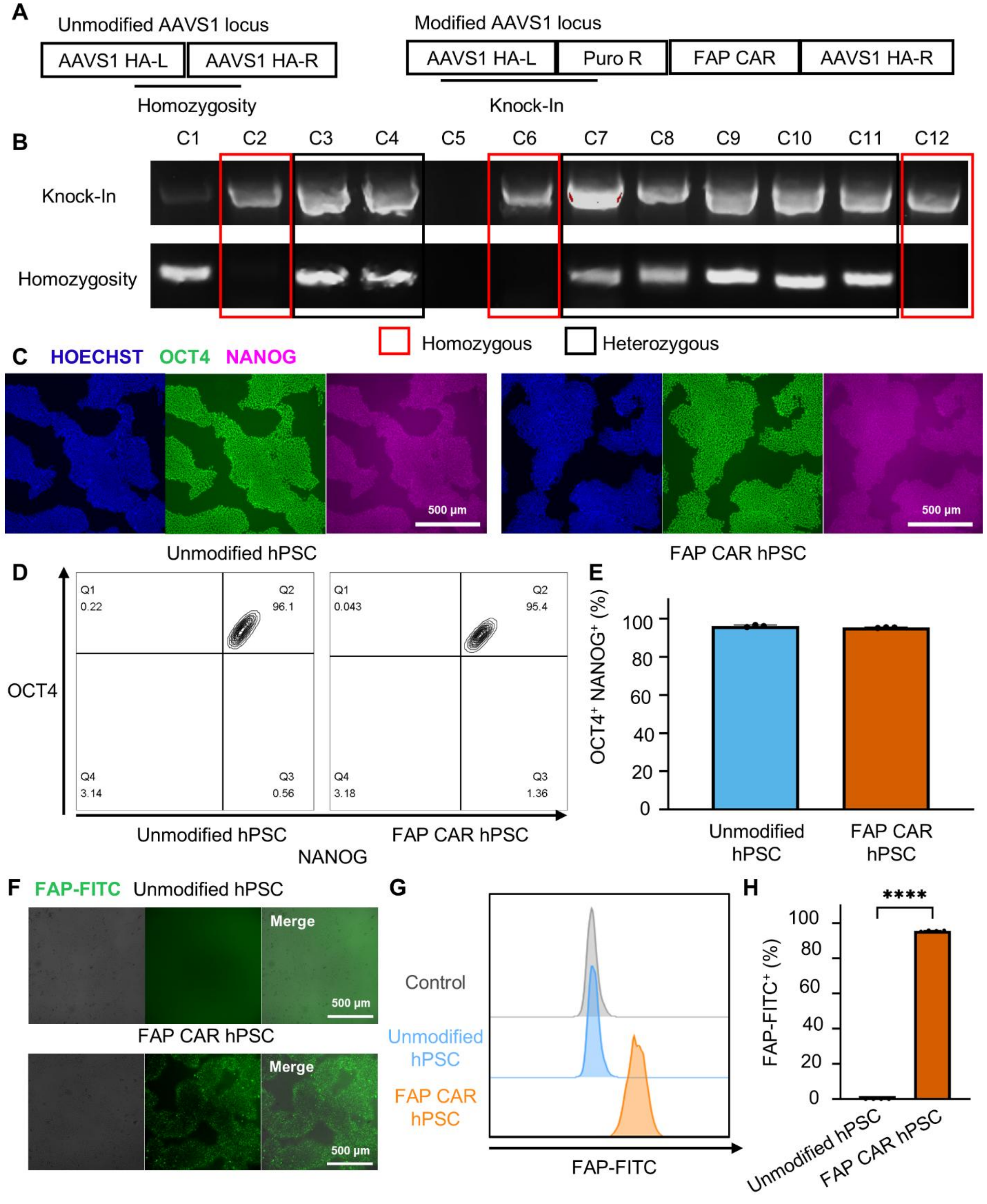


**Supplementary Figure 1. Generation and characterization of FAP CAR hPSCs.**

(A) Schematic detailing the AAVS1 safe-harbor knock-in strategy. (B) Junction-PCR genotyping of edited clones (C1–C12), identifying three homozygous and seven heterozygous clones. (C)

Representative fluorescence images of wild-type and FAP CAR hPSCs expressing the pluripotency markers OCT4 and NANOG. Scale bars, 500 µm. (D) Representative flow cytometry contour plots and (E) quantification of OCT4+ and NANOG+ double-positive cells in wild-type and FAP CAR hPSCs. (F) Representative live-cell images demonstrating FAP-FITC binding on unmodified and FAP CAR hPSCs, indicating surface expression of the FAP CAR. (G) Representative flow cytometry histograms and (H) quantification of FAP-FITC+ cells. Data are represented as mean ± SD. Statistical significance was determined by unpaired Student's t-tests comparing unmodified vs. FAP CAR hPSCs (n = 3 or 4 biological replicates)

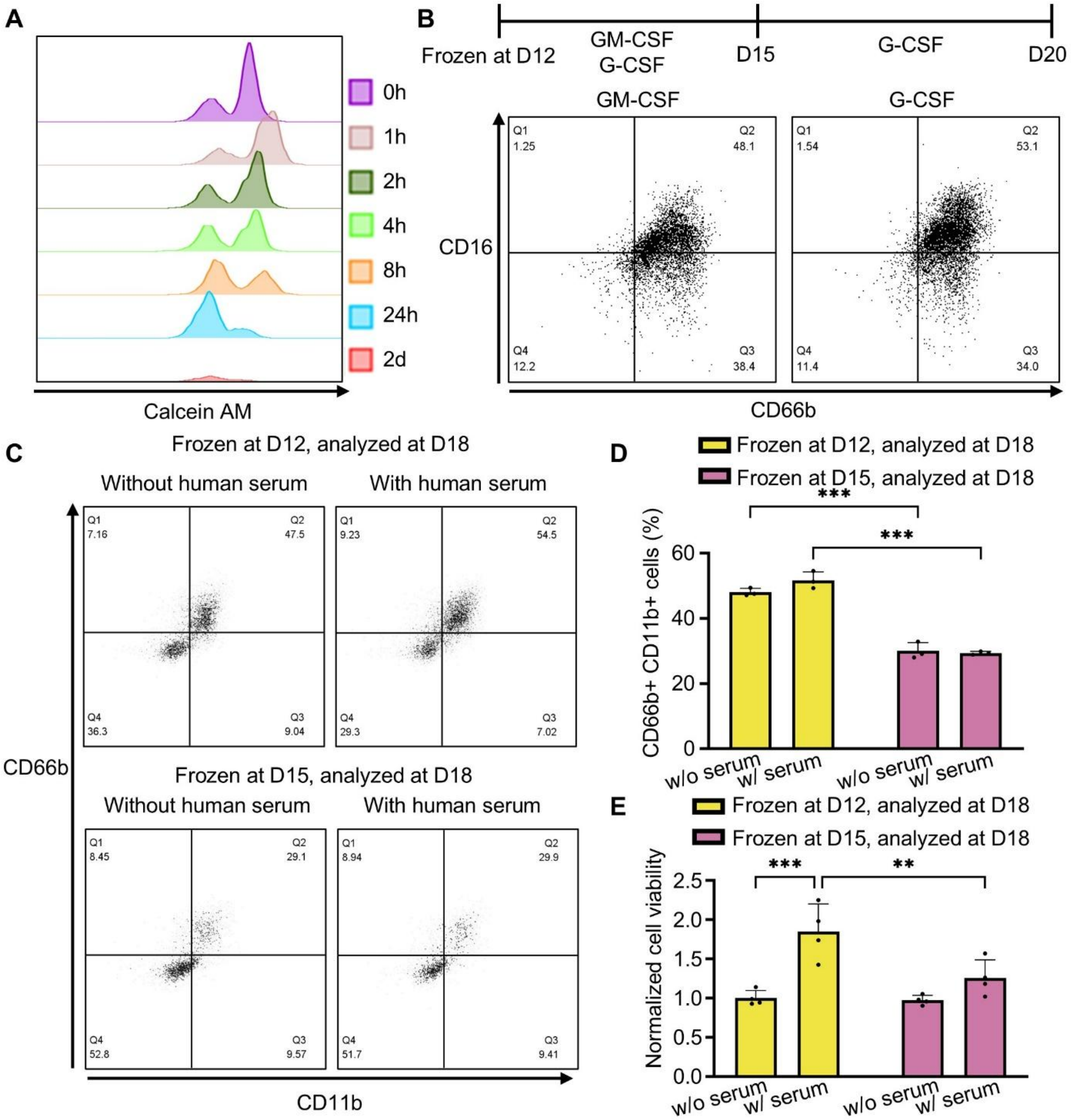

**Supplementary Figure 2. Optimization of cryopreservation conditions for hPSC-derived neutrophil (hNeu) progenitors.**

(A) Post-thaw viability of hPSC-derived neutrophils (hNeu) assessed by Calcein AM staining and flow cytometry analysis at indicated time points (0 h to 2 d) during post-thaw culture. (B) Differentiation efficiency comparison between GM-CSF and G-CSF treatment from day 12 to day 15 using D12-cryopreserved progenitors, assessed by CD16 and CD66b flow cytometry. (C) Representative flow cytometry plots showing CD66b and CD11b expression in progenitors cryopreserved at D12 or D15 (analyzed at D18 and D21, respectively), cultured with or without human serum. (D) Quantification of $CD66b^{+}$ $CD11b^{+}$ double-positive cells from (C) (n = 3). (E) Relative cell viability assessed by CellTiter-Glo 2.0 assay (n = 4). Data are presented as mean ± SEM; statistical significance was evaluated by two-way repeated measures ANOVA (*P < 0.05, **P < 0.01, ***P < 0.001).

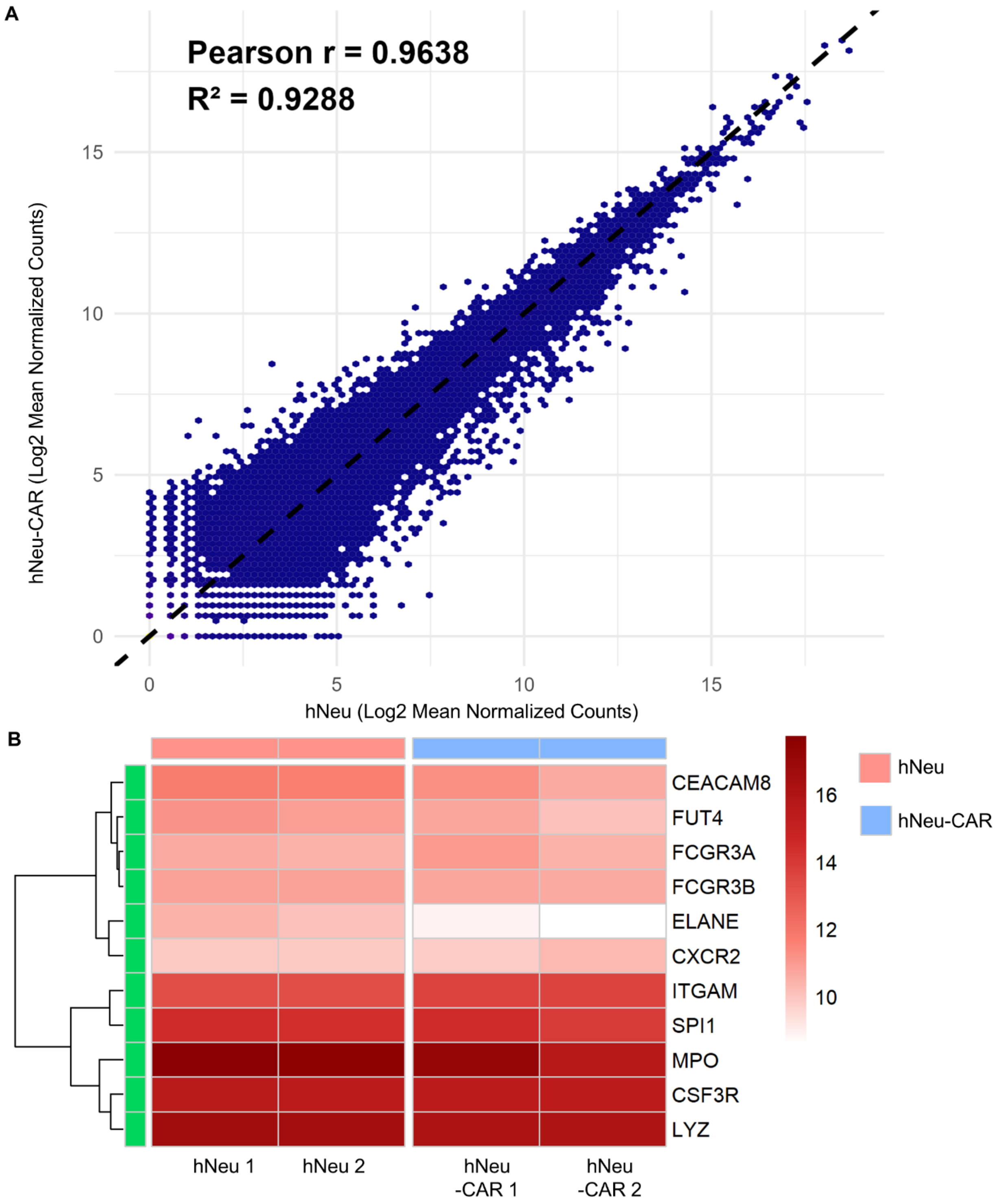


**Supplementary Figure 3. Transcriptional preservation of baseline neutrophil identity in hNeu-CAR cells.**

(A) Global transcriptome correlation scatter plot comparing the mean log2 normalized counts of Control (hNeu) versus CAR hPSC-neutrophils (hNeu-CAR). Pearson r = 0.9638 and R^2 = 0.9288.

(B) Heatmap displaying the expression levels of canonical neutrophil lineage regulators and identity markers in unmodified Control (hNeu 1, 2) and CAR hPSC-neutrophils (hNeu-CAR 1, 2) across two biological replicates.

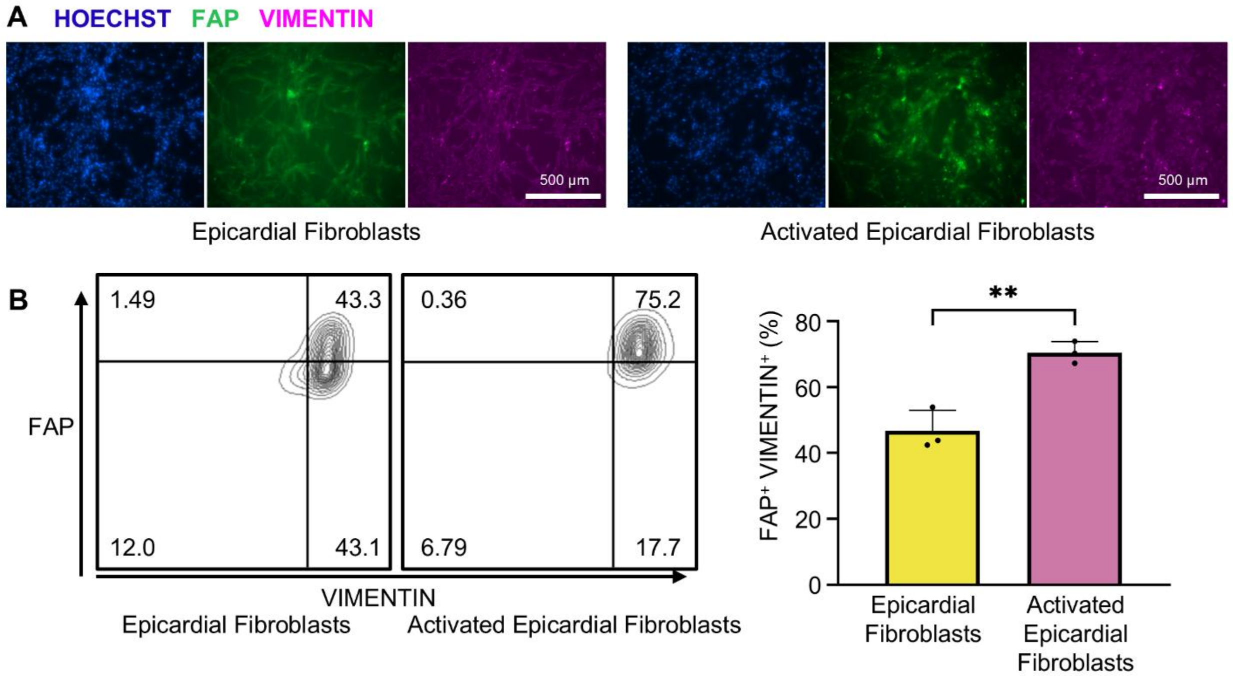


**Supplementary Figure 4. Differentiation and activation of epicardial-derived fibroblasts from hPSCs.**

(A) IF images demonstrating the expression of FAP and VIMENTIN in epicardial-derived fibroblasts, and in fibroblasts activated with TGF-β and FBS. Scale bars, 100 µm. (B) Flow cytometry contour plots and corresponding quantification of FAP+/VIMENTIN+ cells, indicating upregulated FAP expression upon activation. Data are represented as mean ± SD. Statistical significance was determined by unpaired Student's t-tests (n = 3 biological replicates). *P < 0.05, **P < 0.01.

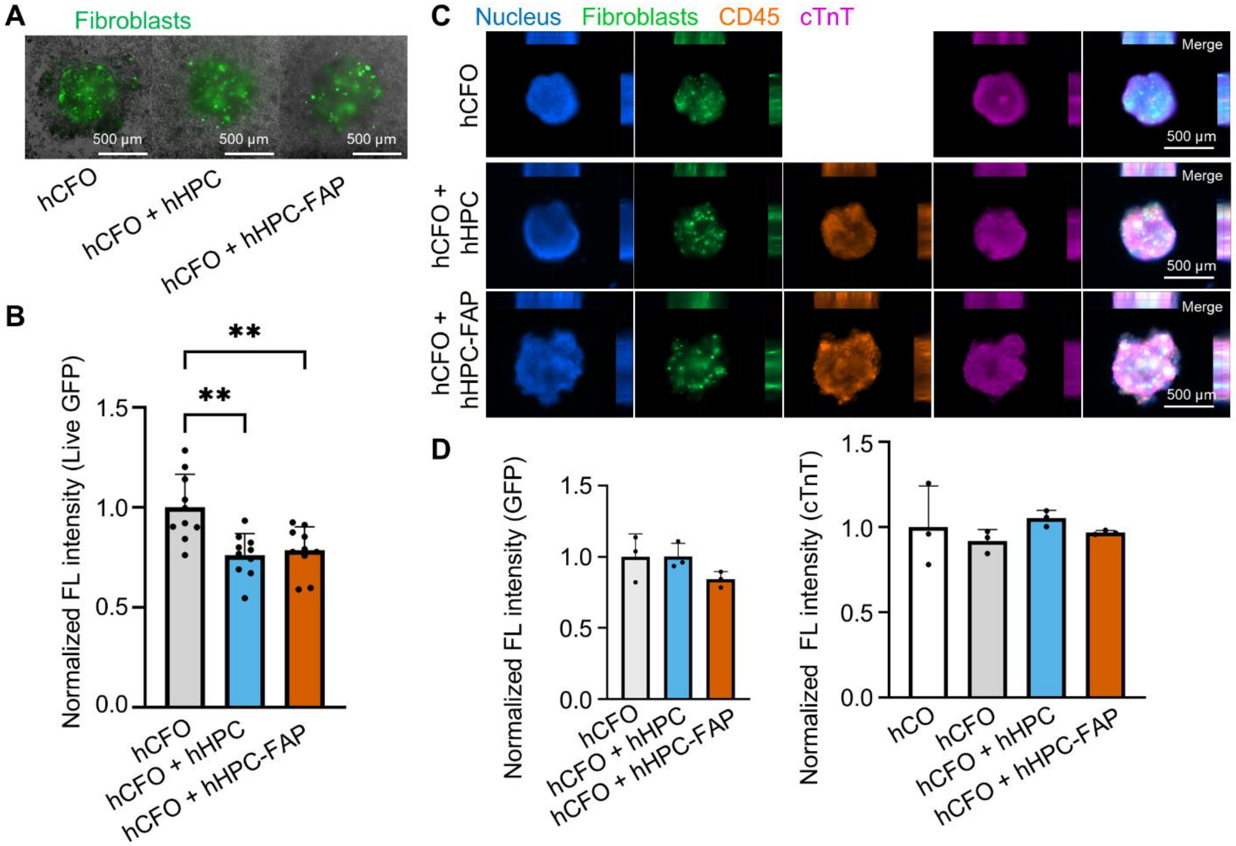


**Supplementary Figure 5. Targeting ability of day 12 hPSC-derived hematopoietic progenitors (hHPCs) and FAP CAR hPSC-derived hematopoietic progenitors (hHPC-FAPs) against activated epicardial fibroblasts in a 3D cardiac organoid model.**

(A) Representative live-cell images of hCFOs containing GFP-expressing epicardial fibroblasts co-cultured with hHPCs or hHPC-FAPs. Scale bars, 500 µm. (B) Quantification of live GFP expression. (C) Representative maximum intensity projections of hCFOs co-cultured with hHPCs or hHPC-FAPs (Green: fibroblasts; Orange: CD45/hHPCs; Magenta: cTnT/CMs). Scale bars, 500 µm. (D) Quantification of GFP (fibroblasts) and cTnT (CMs) fluorescence intensity in co-cultured hCFOs. Data are represented as mean ± SD. n = 10 biological replicates for (B) and n = 3 biological replicates for (D)

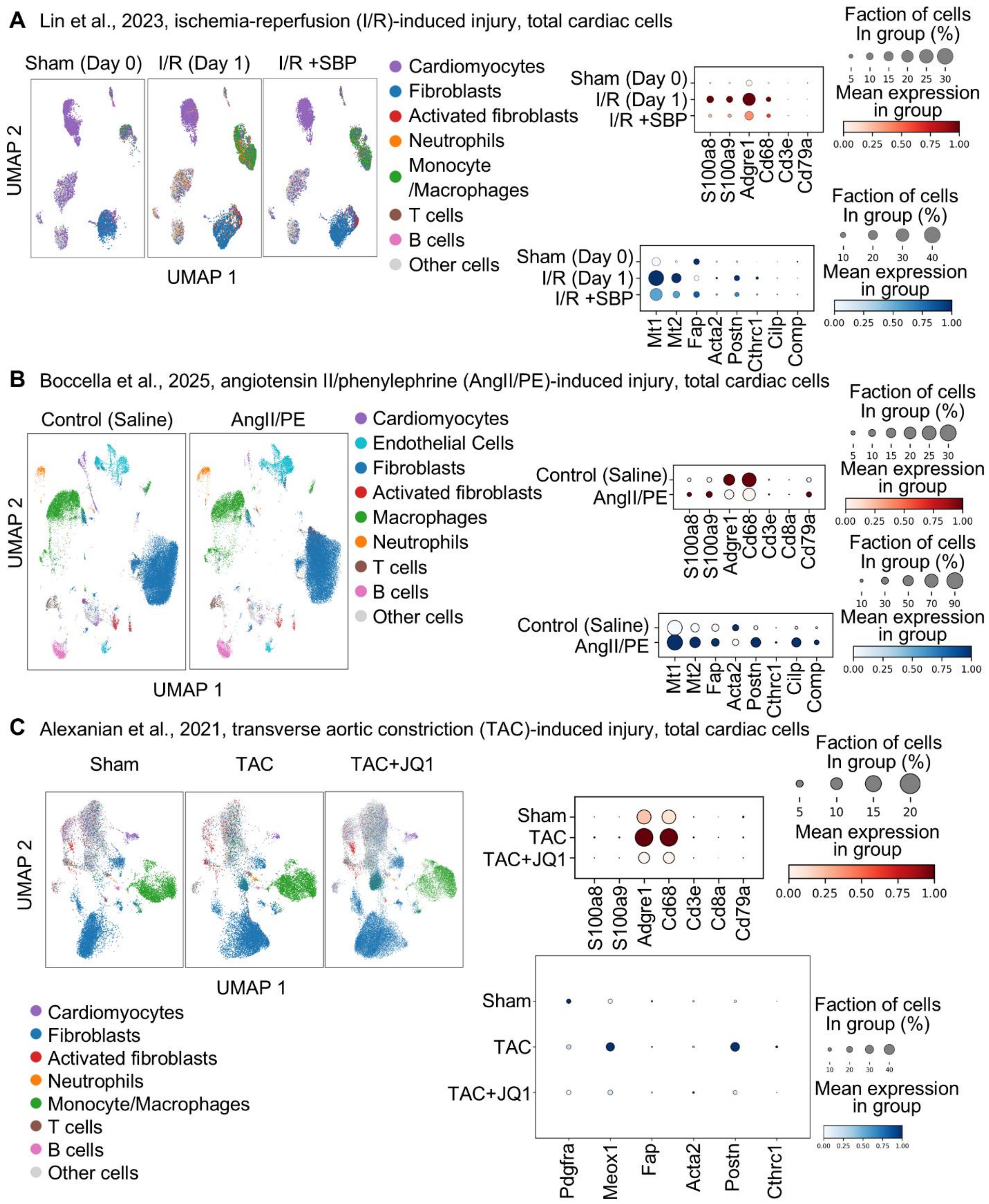


**Supplementary Figure 6. Timing of neutrophil infiltration, immune cell activation, and fibroblast activation in various murine cardiac injury models analyzed via public scRNA-seq datasets.**

(A) Left: UMAP projection of total cardiac cells in a murine ischemia-reperfusion (I/R) model across Sham (Day 0), acute I/R (Day 1), and SBP-treated cohorts (Lin et al., 2023). Right: Dot plots showing the rapid peak of neutrophil (S100a8, S100a9) and myeloid (Adgre1, Cd68) infiltration alongside early stromal stress (Mt1, Mt2) and fibrogenic initiation (Fap, Postn). (B) Left: UMAP projection of whole cardiac cellulome dynamics under continuous neurohormonal stress induced by angiotensin II and phenylephrine (AngII/PE; Boccella et al., 2025). Right: Corresponding dot plots illustrating sustained macrophage dominance and active myofibroblast differentiation (Acta2, Postn, Cthrc1). (C) Left: UMAP embedding of myocardial cells following transverse aortic constriction (TAC)-induced pressure overload with or without BET bromodomain inhibition (TAC+JQ1; Alexanian et al., 2021). Right: Dot plots tracking immune mobilization and the suppression of the profibrotic transcriptional switch (Meox1, Fap, Postn, Cthrc1) under therapeutic intervention. For all dot plots, point size represents the percentage of positive cells within the condition, and color gradients reflect row-scaled mean expression.

**Supplementary Video 1.** Co-culture of $GFP^+$ activated epicardial fibroblasts and hPSC-derived FAP CAR neutrophils at a 1:10 ratio over 24 h. Images were captured every 5 min at 10× magnification using an incubation stage (37 °C, 5% $CO_2$).

**Supplementary Video 2.** Magnified view of Supplementary Video 1 highlighting individual cell–cell interactions.
